# Nonlinear Effects of Intraspecific Competition Alter Landscape-Wide Upscaling of Ecosystem Function

**DOI:** 10.1101/470591

**Authors:** Chelsea J. Little, Emanuel A. Fronhofer, Florian Altermatt

## Abstract

A major focus of ecology is to understand and predict ecosystem function across scales. Many ecosystem functions are only measured at local scales, while their effects occur at a landscape level. Here, we investigate how landscape-scale predictions of ecosystem function depend on intraspecific competition, a fine-scale process, by manipulating intraspecific density of shredding macroinvertebrates and examining effects on leaf litter decomposition, a key function in freshwater ecosystems. Across two species, we found that leaf processing rates declined with increasing density following a negative exponential function, likely due to interference competition. To demonstrate consequences of this nonlinearity, we upscaled estimates of leaf litter processing from shredder abundance surveys in 10 replicated headwater streams. In accordance with Jensen’s inequality, applying density-dependent consumption rates reduced estimates of catchment-scale leaf consumption up to 60-fold versus using density-independent rates. Density-dependent consumption estimates aligned closely with metabolic requirements in catchments with large, but not small, shredder populations. Importantly, shredder abundance was not limited by leaf litter availability and catchment-level leaf litter supply was much higher than estimated consumption, thus leaf litter processing was not limited by resource supply. Our work highlights the need for upscaling which accounts for intraspecific interactions.

## Introduction

In an era of ongoing global change, a growing focus of ecology is to understand what controls ecosystem functioning, and to predict future scenarios of ecosystem function and services. Biodiversity is an important determinant of ecosystem functioning: the traits and dynamics of individuals and species interact to determine the flow of energy and resources through an ecosystem (Hines and Gessner 2012; Grace et al. 2016; Moore and Olden 2017; O’Connor et al. 2017). The number of species in a community is an important determinant of its functioning not just by summing these traits and functions, but also through the interaction between organisms, which can be synergistic or antagonistic (Downing and Leibold 2002; Carrara et al. 2015). Thus, the relative abundance of species is important because the presence of common or dominant species, for example, can influence the relationship between biodiversity and ecosystem function (Smith and Knapp 2003; Dangles and Malmqvist 2004; Winfree et al. 2015). Relative abundance and community assembly are increasingly being incorporated into biodiversity-ecosystem function (BEF) frameworks (for example, Bannar-Martin et al. 2018) as well as being implicitly included in metrics such as evenness and functional diversity, which are sometimes associated with ecosystem function (Hillebrand et al. 2008).

The BEF framework has primarily focused on interspecific interactions. However, intraspecific density-dependent interactions are also recognized as important in almost all disciplines of ecology. For example, they are a key requirement for the maintenance of biodiversity according to modern coexistence theory (Chesson 2000; Amarasekare 2003; McPeek 2012), and many aspects of population dynamics are controlled by density (Hassell et al. 1976; Brook and Bradshaw 2006). Yet intraspecific competition and other density-dependent dynamics rarely appear in BEF schemes partitioning the contribution of different species in a community to ecosystem function (however, see Parain et al. 2018). This is a surprising gap because variation in intraspecific density is ubiquitous in nature: there is considerable variation in species abundances through space and time (Hanski 1990).

Thus, density-dependent control of ecosystem function is not just potentially substantial in magnitude, but could be widespread. The consequences of this mechanism for landscape-level ecosystem functioning remain unexplored, particularly with regard to understanding the shape of a density-ecosystem function (henceforth DEF). Non-linear relationships abound in nature and affect the accuracy of upscaling (Harvey 2000), yet nonlinearities arising from intraspecific interactions are not currently taken into account. Analogous to functional- and temperature-response curves, non-linear DEF relationships would mean that upscaling based on knowledge of ecosystem function at one scale could greatly over- or under-predict gross rates at a broader scale. This idea is captured by Jensen’s inequality, which shows that for nonlinear functions, the mean value of the function across a set of *x* values is not equivalent to the value of the function at the mean of *x* (Jensen 1906), a property which has important consequences for interpreting ecological data (Ruel and Ayres 1999; Martin and Huey 2008; Kingsolver 2009; Denny and Benedetti-Cecchi 2012).

The DEF relationship may be most relevant in determining ecosystem functioning in cases where dominant or highly abundant species contribute a large part of a specific ecosystem function. Widely known and large effect-size examples include salmon importing and exporting nutrients to watersheds (Rüegg et al. 2011) and *Daphnia* clearing lakes of phytoplankton, contributing to secondary production and controlling ecosystem metabolism (Winder and Schindler 2004; Birtel and Matthews 2016). In these cases, a large part of ecosystem function could be predicted by understanding the dynamics of these key taxa, without also considering the comparatively small contributions of rarer taxa. Furthermore, species richness is typically lower where a species is dominant both locally and regionally (Hillebrand et al. 2008), meaning that even in local patches where the dominant species is absent, there may be few other species to provide the same function. For example, in a French stream network, the dominant shredding macroinvertebrate declined in abundance with agricultural intensity, but even where it was absent there were no other taxa which could replace its function in the decomposition process (Piscart et al. 2009). Thus, spatial insurance effects often associated with species turnover (Yachi and Loreau 1999; Loreau et al. 2003) were not present in a way that could maintain this ecosystem function.

In fact, decomposition in freshwater ecosystems may be an ideal setting to explore DEF relationships. Decomposition regulates resource cycling, and is particularly important in aquatic systems where terrestrial detritus can make up a large portion of resource fluxes (Gounand et al. 2018). Furthermore, the characteristic spatial structure of stream networks and the way in which aquatic organisms are limited to dispersing through a stepping-stone arrangement of habitat patches can limit community assembly (Drakou et al. 2009; Brown et al. 2011; Altermatt 2013; Sarremejane et al. 2017; Little and Altermatt 2018*a*). Perhaps partly as a result of this, communities of species contributing to decomposition are characteristically less complex in freshwater than terrestrial ecosystems (Hieber and Gessner 2002), and as a result density variation in those few species could have a large impact (Jonsson and Malmqvist 2003; Klemmer et al. 2012). Decomposition (both in aquatic and terrestrial systems) is less frequently considered in ecosystem function frameworks than is terrestrial biomass production (Cardinale et al. 2011), which may partly explain why DEF has received relatively little attention: for terrestrial producers, a high diversity of species contributes to ecosystem function through time.

Here, we investigate the relationship between intraspecific density of two aquatic macroinvertebrate shredders and their rate of leaf litter processing. To illustrate the potential importance of a nonlinear DEF relationship, we then use these DEF functions to upscale leaf litter processing estimates to catchment levels. Our upscaling exercise is based on spatial variance in intraspecific density of shredders and abundance of leaf litter observed across 10 independent headwater stream networks. Previous upscaled estimates of the effect of shredder species turnover on ecosystem function assumed density-independence of leaf processing rates and furthermore assumed equal densities throughout a catchment (e.g., Piscart et al. 2011). By contrast, we incorporate spatial variance in shredder abundance, and examine qualitative differences in results from upscaling scenarios with and without this spatial variance, for two reasons. First, a previous meta-analysis of laboratory studies indicated density-dependence in per-capita leaf consumption rates (Little and Altermatt 2018*b*). Secondly, in this group of shredding macroinvertebrates typically one species dominates locally, but the dominant species varies in abundance over orders of magnitude within a catchment (Welton 1979; Van den Brink et al. 1991; Altermatt et al. 2016; Little and Altermatt 2018*a*). This creates an ideal scenario to test the concept of a DEF relationship whereby the abundance of these key taxa, rather than species richness, could control decomposition and thus would need to be considered in upscaling predictions.

## Methods

### Study Organisms

We experimentally assessed effects of intraspecific density on leaf shredding rates by two freshwater amphipod (Crustacea, Amphipoda) species: *Gammarus fossarum* (Koch), a relatively small species native to Central Europe, and *Dikerogammarus villosus* (Sowinsky), a relatively large species native to the Ponto-Caspian region which has recently invaded many regions worldwide (Van den Brink et al. 1991; Gallardo and Aldridge 2015; Šidagytė et al. 2017). As a guild, amphipods are the dominant macroinvertebrates, including the dominant invertebrates in the shredding functional group, in many central European streams (for examples, Piscart et al. 2009; Nery and Schmera 2015). Collection and maintenance of study organisms are described in the Supplementary Material.

### Mesocosm Experiments

Mesocosms were built from 2 L plastic containers with 0.4 m^2^ of bottom surface area, placed in a flowing-water rack system with a mixture of stream and tap water. Conditioned naturally senescent alder leaves totalling 1.5 g (dry weight) were placed in each mesocosm. Alder is commonly found in benthic leaf litter samples in headwater streams in this area (Little and Altermatt 2018*c*) and is a preferred food source for these species (Little and Altermatt 2018*b*). For each species, mesocosms were set up with fixed densities of the target amphipod species: 50 replicates with one individual, 20 replicates with two individuals, 10 with five individuals, 10 with 10 individuals, six with 20 individuals, and six with 30 individuals per mesocosm. This 30-fold density range is smaller than the >100-fold density range commonly observed in stream reaches (Little and Altermatt 2018*a*). The unbalanced number of replicates for each density was chosen because per-mesocosm leaf consumption was expected to be more variable in replicates with fewer amphipods.

The leaf consumption experiments were run for 19 (*G. fossarum*) and 12 (*D. villosus*) days, respectively, at which point leaves from the mesocosms were collected and dried for 48 h at 60 °C, then weighed to calculate mass loss from the beginning of the experiment. Resources remained *ad libitum* throughout the experiment, and at least 0.65 and 0.55 g of leaf litter remained at the end of the experiment for *G. fossarum* and *D. villosus*, respectively (representing ≥ 43% and ≥ 36% of the resources initially available). Amphipods were counted every two to three days throughout the experiments to track mortality; overall, survival was 89.3% for *G. fossarum* and 95.1% for *D. villosus*. These mortality estimates were used to calculate an average amphipod density (individuals per square meter) that the mesocosm experienced over the length of the experiment. At the end of the experiment, amphipods were sacrificed, dried for 48 h at 60 °C, and weighed. Individuals which died during the course of the experiment were assigned the global average weight of all amphipods across the experiment. The average daily biomass in a mesocosm (mg m^−2^) was then calculated as the average density multiplied by the average weight of all individuals in the mesocosm. Two outliers were removed from the *G. fossarum* dataset and three from the *D. villosus* dataset, because their consumption rate estimates were over three standard deviations from the mean and also substantially higher than any we had measured in previous experiments (Little and Altermatt 2018*b*), and we did not feel we could rule out measurement error as an explanation.

### DEF models

For both amphipod species, we tested for the effects of density on leaf consumption using nonlinear models in R version 3.5.0 (R Core Team, Vienna, Austria). Initial data exploration and linear models using transformed and non-transformed data showed that these relationships were linear in log-log space (see Supplementary Material for details, and Figures S1-S3). Therefore we created negative exponential models using the gNLS function in the ‘nlme’ package version 3.1-137 (Pinheiro et al. 2013) and weighted data points by the variance in the response variable, since there was higher variance around high estimates of leaf litter consumption across the experiment. For each species we created separate models of the relationships between amphipod density and per-amphipod daily leaf consumption, and between amphipod biomass and biomass-adjusted daily leaf consumption.

### Field Surveys and Upscaling of Shredder Abundance

We upscaled estimates of leaf litter processing to the catchment level by pairing the derived DEF equations with spatially resolved population density data from field surveys. We had previously assessed amphipod abundance in ten headwater stream catchments in eastern Switzerland predominantly inhabited by *G. fossarum*, where *D. villosus* was present only rarely at the outlets (Altermatt et al. 2016; Little and Altermatt 2018*a*); the latter species is more common in rivers (Van den Brink et al. 1991). The goal of upscaling was to demonstrate the consequences of nonlinear DEF relationships, so since shape of the relationship was similar in the two species, we performed the analysis based only on the *G. fossarum* DEF function. The full details of the field surveys are described in Little and Altermatt (2018*b*), but briefly, sampling points were established in April 2015 in every ∼250 m section of each stream. Amphipods were collected using a kicknet and their density was estimated on a logarithmic scale (0, 1–10, 11–100, 101–1000, or >1000 individuals per 1 meter-long stream segment). Below, we refer to these abundance estimates as ‘bins’.

For upscaling, we longitudinally divided each stream’s mapped watercourse (Swisstopo 2007) into one-meter segments and used two different methods to estimate the total abundance of amphipods in the catchment: inverse distance weighted interpolation and proportional estimation. We simulated spatial abundances of amphipods 1000 times per catchment using each method, and averaged over the simulations to extract catchment-wide predictions of abundance and processing (see below).

Inverse distance weighting (IDW) produces interpolated data that varies smoothly in space as a function of distance from measured sampling points, based on the assumption that points close to each other are more similar. Each IDW simulation began by assigning the catchment’s sampling points (n = 9–15, depending on the catchment) to a random abundance value within their observed abundance bin (e.g., a random number between 11 to 100 for a bin with 11–100 individuals). Then using the package ‘gstat’ version 1.1-6 (Pebesma 2004), each one-meter segment was assigned an abundance based on its distance from these 9–15 assigned points.

With the proportional estimation method, we removed the assumption that nearby reaches are more similar to each other and instead focused only on capturing the observed variation in surveyed abundances. With this method, we recorded the proportion of a catchment’s sampling points which belonged to each abundance bin (i.e., proportion of sampling points with zero amphipods, proportion with 1–10 amphipods, etc.) and created a probability distribution of abundance bin assignment for the catchment. For a simulation, every one-meter stream segment was randomly assigned to an abundance bin based on this probability function, and then the segments were assigned random abundances from within the range of their assigned bins (e.g., assigned to the 1–10 amphipod bin, and then assigned a random number between 1 and 10).

For each simulation, the one-meter segments were summed to produce a catchment-level abundance estimate. The 1000 simulations per catchment (per method) were summarized with means and 95% confidence intervals.

### Upscaling Processing Rates to Real Catchments

We estimated whole catchments’ total leaf litter processing rate per day based on these abundance estimates, under two scenarios. In Scenario 1, we multiplied the global average per-capita processing rate from the *G. fossarum* density experiment (i.e., average across all densities) by the total population size of the catchment, a common way to upscale consumption estimates (for example, Piscart et al. 2011). In Scenario 2, we used the spatially-varying amphipod densities derived from the two estimation methods, and applied the experimentally-derived *G. fossarum* DEF function from to each one-meter stream segment before summing to the catchment level.

To compare these estimates to another frame of reference for understanding organisms’ food consumption rates, we also estimated catchment-level leaf litter consumption rates based on metabolic requirements. We converted the total estimated abundance of *G. fossarum* in each catchment to biomass by multiplying their number by the average dry mass of *G. fossarum* used in the mesocosm experiments, 2.85 mg. We then estimated the amount of leaf litter consumption required by this biomass of amphipods on an annual scale, using three steps. (1) The production:biomass ratio (P/B) was set at 3.5 as in prior woodland stream macroinvertebrate work (Petersen et al. 1989); this is intermediate between P/B estimates of congeneric taxa, including 4.29 for lake amphipods (Zhang et al. 2016), 4.65 for *Gammarus pseudolimnaeus* in a river (Marchant and Hynes 1981), and 2.6 for *Gammarus pulex* in a stream (Mortensen 1982). (2) Resources are required both for production and basic metabolic maintenance; as such, the ratio of respiration:production for macroinvertebrates has been estimated at 70:30 (Cummins 1975), such that 0.3 is the production:assimilation ratio (P/A). (3) Finally, *D. villosus* and another gammarid amphipod, *Gammarus roeselii*, were found to have identical 40% assimilation efficiencies of conditioned leaves (Gergs and Rothhaupt 2008), so we used a 0.4 assimilation:consumption ratio (A/C). Thus, per mg of amphipod biomass, we calculated:

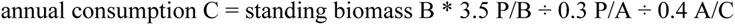

This produced a consumption:biomass ratio of 29.2, similar to the ratio of 22.7 used for amphipods and isopods in the Lake Erie ECOPATH model (Zhang et al. 2016). We did not use this value directly because lakes contain different amphipod species than headwater streams. Furthermore, while both groups of amphipods are omnivores with flexible feeding strategies, lake amphipods feed substantially on diatoms, phyto- and zooplankton (Covich et al. 2010), which have different nutritional content than leaf litter; it is reasonable that the group consuming lower-quality food sources should need more of it to meet metabolic requirements.

### Leaf Litter Availability in Real Catchments

In parallel to assessing amphipod densities for upscaling processing rates, we also assessed the availability (abundance) of leaf litter in the stream catchments at the same spatio-temporal resolution. (Little and Altermatt 2018*c*). This allowed us to put the processing rates into context, and also could be used determine whether processing of leaves was donor-limited (i.e., whether leaf litter was consistently available as a food resource). We used two methods to assess leaf litter supply and compare it to the estimated demands from the upscaling estimates. First, at the same sampling points in the ten catchments where we surveyed amphipod abundance, we concurrently measured benthic leaf litter standing stock four times over the course of a year (Little and Altermatt 2018*c*). Briefly, at each sampling point and at each sampling visit, the substrate of the 1 m long stream segment was classified into substrate types and vegetation and benthic leaf litter cover using a 1 × 1 m sampling frame with 0.2 × 0.2 m gridlines (further details in Little and Altermatt 2018*b* and *c*). Then, all benthic leaf litter was collected from a known subsample of area (mean area collected = 0.032 m^2^ ± 0.012 m^2^ s.d.), and the substrate area of the subsample as a proportion of the total amount of streambed covered in leaf litter in that stream segment was used to calculate the standing stock of leaf litter (in g dry weight m^−2^) at that sampling point. We a priori hypothesized that amphipod distributions would be correlated with leaf litter availability, however this was only minimally borne out in a joint species distribution model based on amphipod presence/absence (Little and Altermatt 2018*a*). To assess whether amphipod abundance was correlated with leaf litter availability, here, we summarized the distribution of benthic leaf litter standing stock measurements for each abundance class of amphipods. We made a linear mixed-effect model using the ‘lme4’ package version 1.1-18-1 (Bates et al. 2015) with benthic leaf litter as a fixed factor and sampling point and season as random factors to account for repeated sampling visits, and square-root transformed minimum of the abundance class as the response variable. The significance of the fixed factor was assessed using a Type III analysis of variance with Satterthwaite’s method with the ‘lmerTest’ package version 3.0-1 (Kuznetsova et al. 2015).

Second, we estimated annual leaf litter input to each of the ten catchments by combining field sampling with land cover data. We deployed leaf litter traps (n = 8 per site, collecting an area of 800 cm^2^ each) at six different sites (3 deciduous forest, 1 mixed forest, 2 agricultural) during fall leaf drop (September 8 to December 1, 2015). We then paired this with data on land cover within a 1 m buffer zone on either side of each stream. This represents a conservative estimate of leaf litter input because we did not measure lateral blow-in, which occurs throughout the year and can represent an additional ∼20–50 % of total leaf litter inputs to forested headwater streams (Fisher and Likens 1973; Conners and Naiman 2008; Kochi et al. 2010). Land cover assessment was primarily based on the 2012 CORINE land cover/land use classification (Bossard et al. 2000), which was produced from Indian Remote Sensing (IRS) P6 LIS III and RapidEye imagery with a Minimal Mapping Unit of 25 hectares and positional accuracy of, at a minimum, 100 m. The CORINE data, however, did not reflect the presence of riparian strip vegetation (either shrubs or trees) within agricultural, suburban, or industrial land use types. To incorporate this information, we manually delineated it based on 25 cm resolution color orthophotos (Swisstopo 2016). The total area in the 1 m buffer zone for each stream catchment was thus summarized into categories of forested, other riparian tree/shrub, or other cover. The forested area in the buffer was assigned the mean input rate from our leaf litter traps deployed in forested areas. The other riparian tree/shrub area was assigned half this input rate, because it was difficult to delineate shrub vs. tree vegetation from orthophotos; the 50% choice is conservative, as shrub-dominated riparian strips can have similar or only slightly lower leaf litter input rates than forests in some cases (Delong and Brusven 1994; Scarsbrook et al. 2001). Other land cover types in the buffer area were assigned zero leaf litter input per year, a third deliberately-conservative assumption in our estimation process. For each catchment, we summed the input from each of these area types to obtain a total, lower-bound estimate of annual leaf litter input.

## Results

Our experimental data fit negative exponential functions relating per-capita consumption rates to density (for fitting details, see Supplementary Material). This was true both when relating individual density to per-capita leaf consumption (Figure 1), and density of biomass to biomass-adjusted leaf consumption (Figure S4). To confirm that these derived relationships explained the density-dependent relationship with per-capita consumption rates, we calculated predicted per-mesocosm total leaf consumption along a continuous gradient of amphipod densities. These curves (solid lines in right panels of Figure 1) reasonably matched the actual per-mesocosm leaf consumption rates, while linear extrapolations based on density-independent, constant per-capita consumption rates overestimated total leaf consumption by orders of magnitude for any density greater than a few amphipods per square meter (Figure 1).

**Figure 1.**
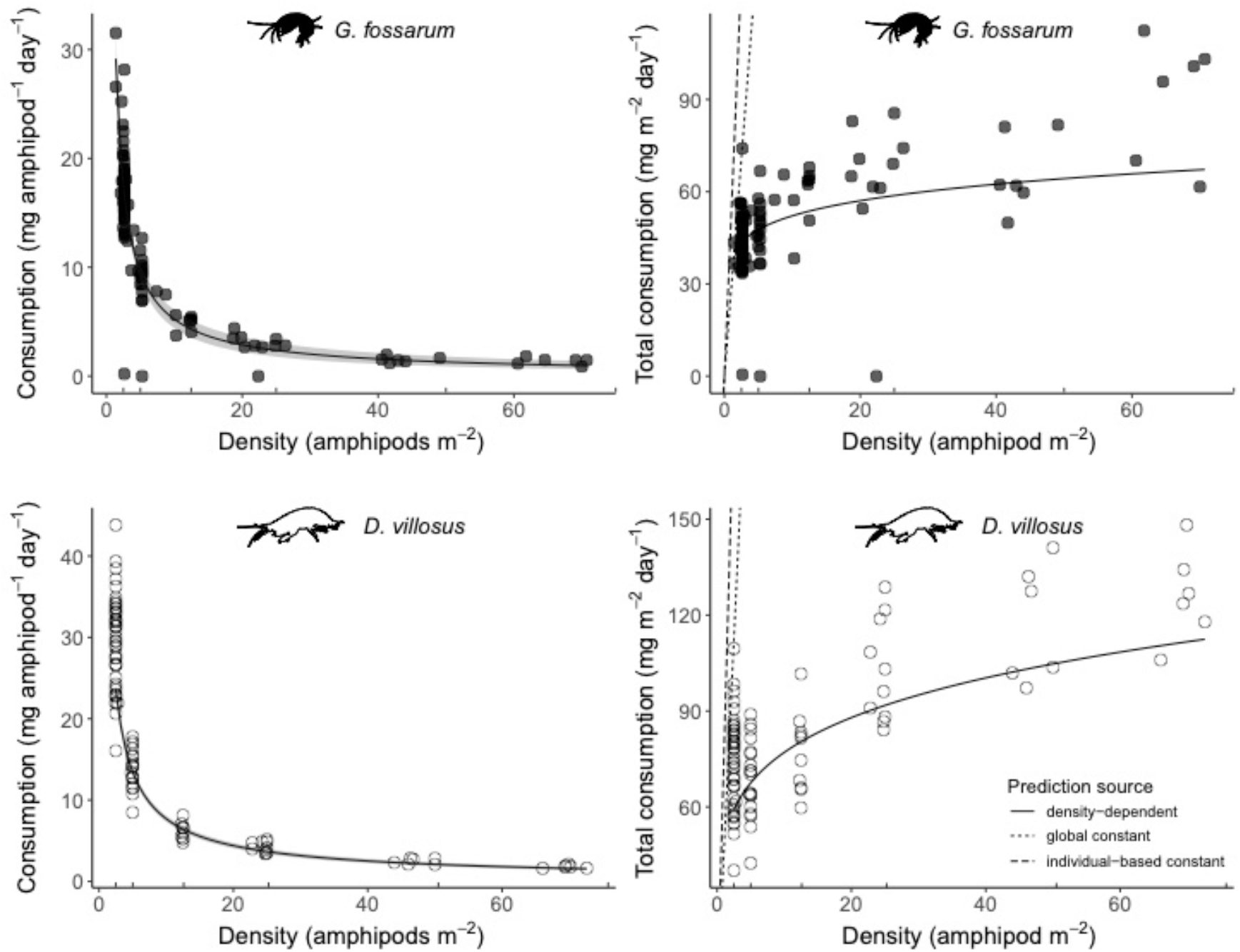
Negative exponential relationships (left panels) between density and per-capita consumption rates for G. fossarum (consumption = 38.6*density-0.87) and D. villosus (consumption = 49.7*density-0.81) in mesocosm experiments. Gray shading shows the 95% confidence interval of the model fit. Right panels show the total daily leaf litter consumption per mesocosm, overlaid by expected values from the negative exponential functions derived at left (solid lines), constant per-capita consumption rates calculated by averaging all mesocosms (dotted line), and constant per-capita consumption rates calculated only from mesocosms with one individual amphipod each (dashed line). Note the different x-and y-axis ranges for the two species. Vertical tickmarks above the x-axis indicate starting densities for experimental mesocosms.

Next, we performed upscaling to estimate amphipod abundance in the ten study catchments. Estimates of whole-catchment abundance ranged from hundreds (808 in Dorfbach) to millions (1.46 million in Mannenbach) of amphipods using the inverse distance weighted estimation method (Table 1), and from thousands (1,590 in Dorfbach) to millions (1.43 million in Seebach) using proportional estimation (Table S2).

**Table 1.** Total amphipod abundance estimated using inverse distance weighted interpolation based on field sampling (*n* = number of sampling points in catchment, *l* = total stream length in the catchment), and three different estimates to whole-catchment leaf litter processing (grams/day). Briefly, the first two scenarios are based on our laboratory data of leaf litter consumption rates: Scenario 1 assumes density-independent per-capita processing, while Scenario 2 assumes spatially-varying abundances and density-dependent leaf consumption calculated for each stream reach. The third estimate is based on metabolic requirement for growth and respiration, assuming 40% assimilation of ingested leaf litter. Estimates are on means of 1000 simulations of total abundance, with 95% confidence intervals shown in parentheses.

| Stream name | Total Abundance | Scenario 1 Processing | Scenario 2 Processing | Metabolic requirement |
| --- | --- | --- | --- | --- |
| Chesselbach<br>$n = 13, l = 4.5 \text{ km}$ | 720,719<br>($\pm 1,035$) | 8,194.7<br>($\pm 122.1$) | 315.4<br>($\pm 0.5$) | 164.1<br>($\pm 0.2$) |
| Dorfbach<br>$n = 15, l = 4.3 \text{ km}$ | 808<br>($\pm 6$) | 9.2<br>($\pm 0.3$) | 9.8<br>($\pm 0.1$) | 0.2<br>($\pm 0.0$) |
| Eschlibach<br>$n = 12, l = 4.1 \text{ km}$ | 1,135,006<br>($\pm 1,249$) | 12,905.2<br>($\pm 145.9$) | 312.6<br>($\pm 0.49$) | 258.5<br>($\pm 0.3$) |
| Hepbach<br>$n = 13, l = 3.8 \text{ km}$ | 231,020<br>($\pm 622$) | 2,626.7<br>($\pm 67.6$) | 178.3<br>($\pm 0.7$) | 52.6<br>($\pm 0.1$) |
| Imbersbach<br>$n = 11, l = 3.3 \text{ km}$ | 139,222<br>($\pm 613$) | 1,583.0<br>($\pm 45.1$) | 119.0<br>($\pm 1.2$) | 31.7<br>($\pm 0.1$) |
| Mannenbach<br>$n = 15, l = 4.8 \text{ km}$ | 1,464,130<br>($\pm 1,408$) | 16,647.4<br>($\pm 178.1$) | 378.0<br>($\pm 0.6$) | 333.4<br>($\pm 0.3$) |
| Seebach<br>$n = 11, l = 3.2 \text{ km}$ | 1,370,115<br>($\pm 1,250$) | 15,578.4<br>($\pm 139.1$) | 260.4<br>($\pm 0.3$) | 312.0<br>($\pm 0.3$) |
| Tobelmühlibach<br>$n = 12, l = 3.7 \text{ km}$ | 696,563<br>($\pm 1,081$) | 7,920.0<br>($\pm 107.8$) | 239.9<br>($\pm 0.5$) | 158.5<br>($\pm 0.2$) |
| Unnamed Stream 1<br>$n = 9, l = 2.8 \text{ km}$ | 518,543<br>($\pm 849$) | 5,895.9<br>($\pm 120.8$) | 172.1<br>($\pm 0.6$) | 118.1<br>($\pm 0.2$) |
| Unnamed Stream 2<br>$n = 10, l = 3.6 \text{ km}$ | 317,053<br>($\pm 672$) | 3,604.9<br>($\pm 64.2$) | 234.3<br>($\pm 0.4$) | 72.2<br>($\pm 0.2$) |

In an example catchment, the Chesselbach (for all other catchments, Table 1, Table S2, and Figures S5–S13), inverse distance weighted interpolation from 13 sampling points (Figure 2) produced an estimate of ∼720,000 amphipods in the catchment (mean of 1000 simulations: 720,719; 95% CI 709,980–731,456). Using the mean experimental per-capita consumption rate (12 mg amphipod^−1^ day^−1^) to derive leaf processing (Scenario 1) yielded a mean of 8.8 kg of leaf litter (dry weight) processed per day. Applying the experimentally-derived negative exponential DEF relationship to the spatially-varying interpolated densities in the catchment (Scenario 2) resulted in a markedly lower predicted processing rate, in this case a mean of 0.3 kg per day. Indeed, in all but one (the most sparsely occupied) catchment, estimates of total leaf litter processing were lower using the experimentally-derived DEF relationship than when using a density-independent processing rate (Table 1). The mismatch was substantial: not accounting for density dependence resulted in leaf processing rates up to 60 times higher in some catchments. Results were similar when based on proportional abundance estimations (Table S2). Because stream reaches with higher densities of individuals have the lowest per-capita processing rates, the spatial distribution of leaf litter processing under this scenario is very different than the spatial distribution of amphipod abundance, with the effect of homogenizing ecosystem function in space despite having heterogeneous biomass (Figure 2). The processing estimates based on Scenario 2 corresponded well with estimated metabolic requirements for amphipod populations of the estimated size (Table 1). In all catchments with total abundance greater than 500,000 amphipods, which corresponded to average densities of more than 100 amphipods per meter of stream length, the metabolic estimate was within 50% of the Scenario 2 estimate. The Scenario 1 estimates, by contrast, were 50 times greater than the estimated metabolic requirements.

**Figure 2.**
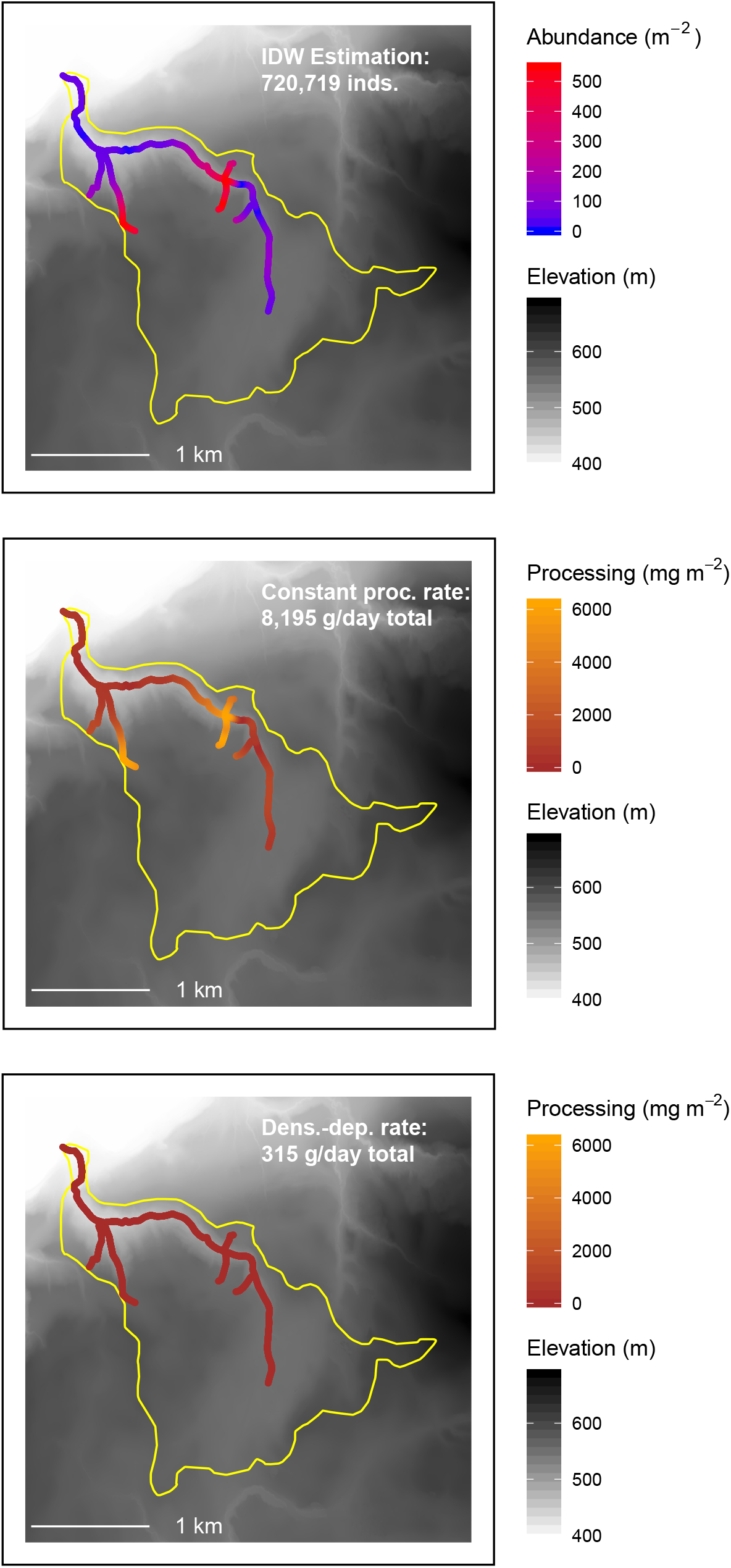
Hotspots of abundance (top panel) and litter processing (bottom panels) in the Chesselbach catchment (yellow outline). Abundance at 13 sampling points was scaled up using inverse distance weighted interpolation. Daily processing rates were calculated by multiplying the interpolated abundance in a 1 m section of stream length by either the average per capita consumption rate of G. fossarum (middle panel) or the experimentally-derived negative exponential function relating *G. fossarum* density to per-capita leaf litter consumption (bottom panel). This figure shows the mean of 1000 simulations of the interpolation process. Data sources: swisstopo (2010, 2014), Vector25 and TLM3D, DV 5704 000 000, reproduced by permission of swisstopo/JA100119.

Greater benthic leaf litter standing stock did not correspond to higher abundances of amphipods (Figure 3): the fixed and random factors combined explained more variance (conditional R^2^ = 0.29) than the fixed factors alone (marginal R^2^ = 0.01), and the fixed factor of leaf litter availability was not significantly associated with amphipod abundance (F_1,402_ = 0.14, p = 0.70). This was congruent with its overall low contribution to variation in amphipod presence/absence when assessed together with other explanatory variables such as water chemistry, land use, and microhabitat (Little and Altermatt 2018*a*). However, low values of standing stock could be due to multiple mechanisms, including low input rates, high processing rates, or flushing downstream. Therefore, we also estimated total leaf litter input rates. From leaf litter traps deployed in forested areas capturing vertical litterfall during the fall leaf drop period, an average of 478 g m^−2^ (dry weight) of litter inputs were available annually (range: 373–509 g m^−2^, n = 4 sites; Figure S14). We then mapped these inputs onto land cover patterns in the catchments. All but four catchments had > 50 % of their 1-meter buffer zone in forested areas, and these four catchments had 53–91 % of their buffer area in riparian strip vegetation (Figure 4A). Overall, non-shrub/tree cover accounted for only 2–28 % of the near-stream buffer zone, except for one outlier catchment (Seebach) for which it represented 47 % of the buffer area (Figure 4A). Seebach thus had the lowest estimated leaf litter inputs (77 kg yr^−1^), while the other catchments received an estimated 1084–3533 kg of leaf litter from their directly adjacent vegetation annually (Figure 4B). For most catchments this represented 18–30 times the amount of leaf litter consumed annually in our upscaling estimates under Scenario 2, which incorporated spatial heterogeneity in abundances as well as density-dependent consumption (Figure 4B). The exceptions were for a sparsely-inhabited but heavily vegetated catchment (Dorfbach), where it was 700 times the estimated consumption requirement, and a densely-inhabited but sparsely-vegetated catchment (Seebach), where it was only eight times the estimated consumption requirement.

**Figure 3.**
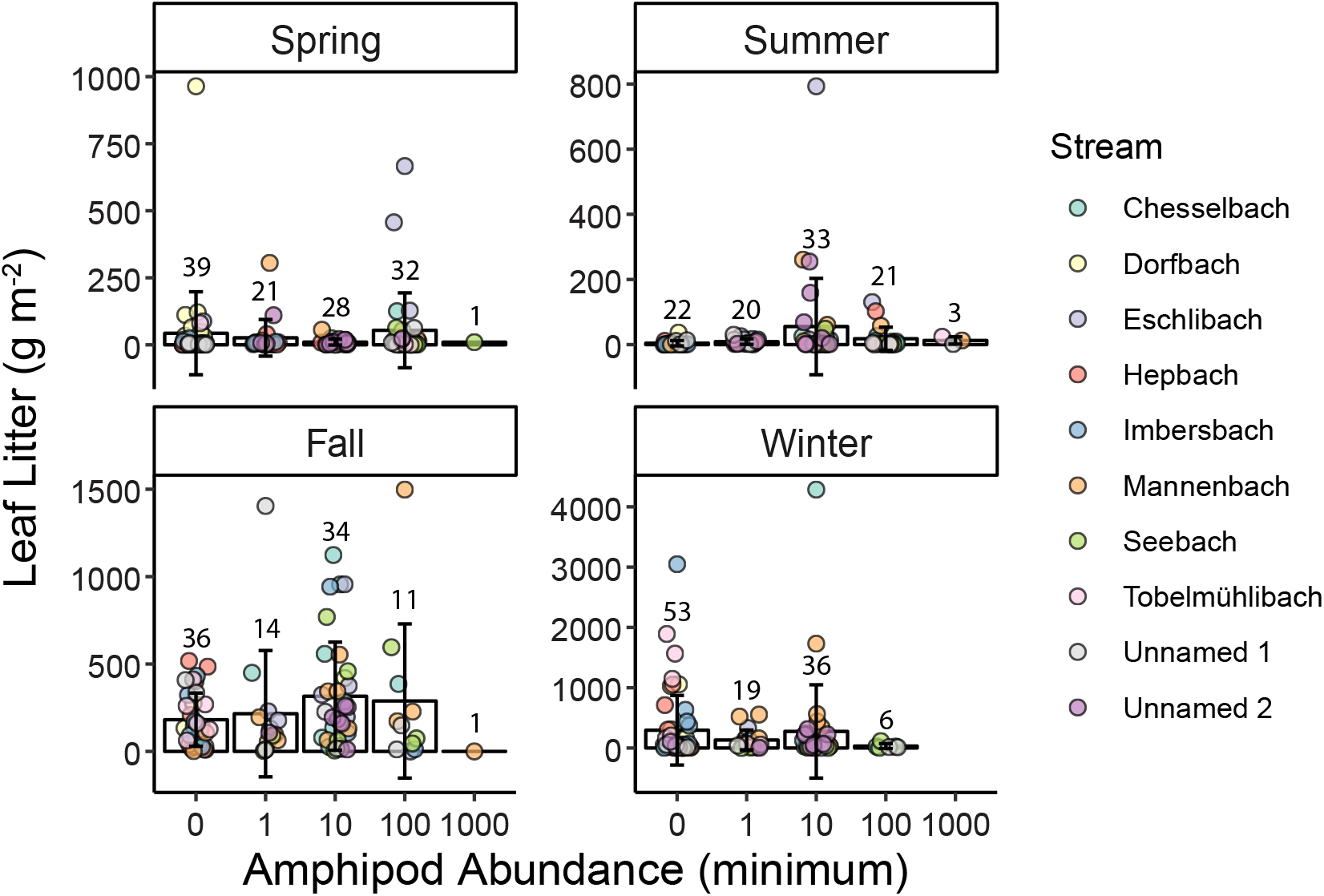
Standing stock of benthic leaf litter at sampling points where different abundance classes of amphipods were observed. Error bars show standard deviation of observations within an abundance class, and colored points show raw values. Numbers above the error bars indicate the number of sampling points falling into each abundance class in each sampling visit. Note that y-axis is transformed, and has a different range for each season.

**Figure 4.**
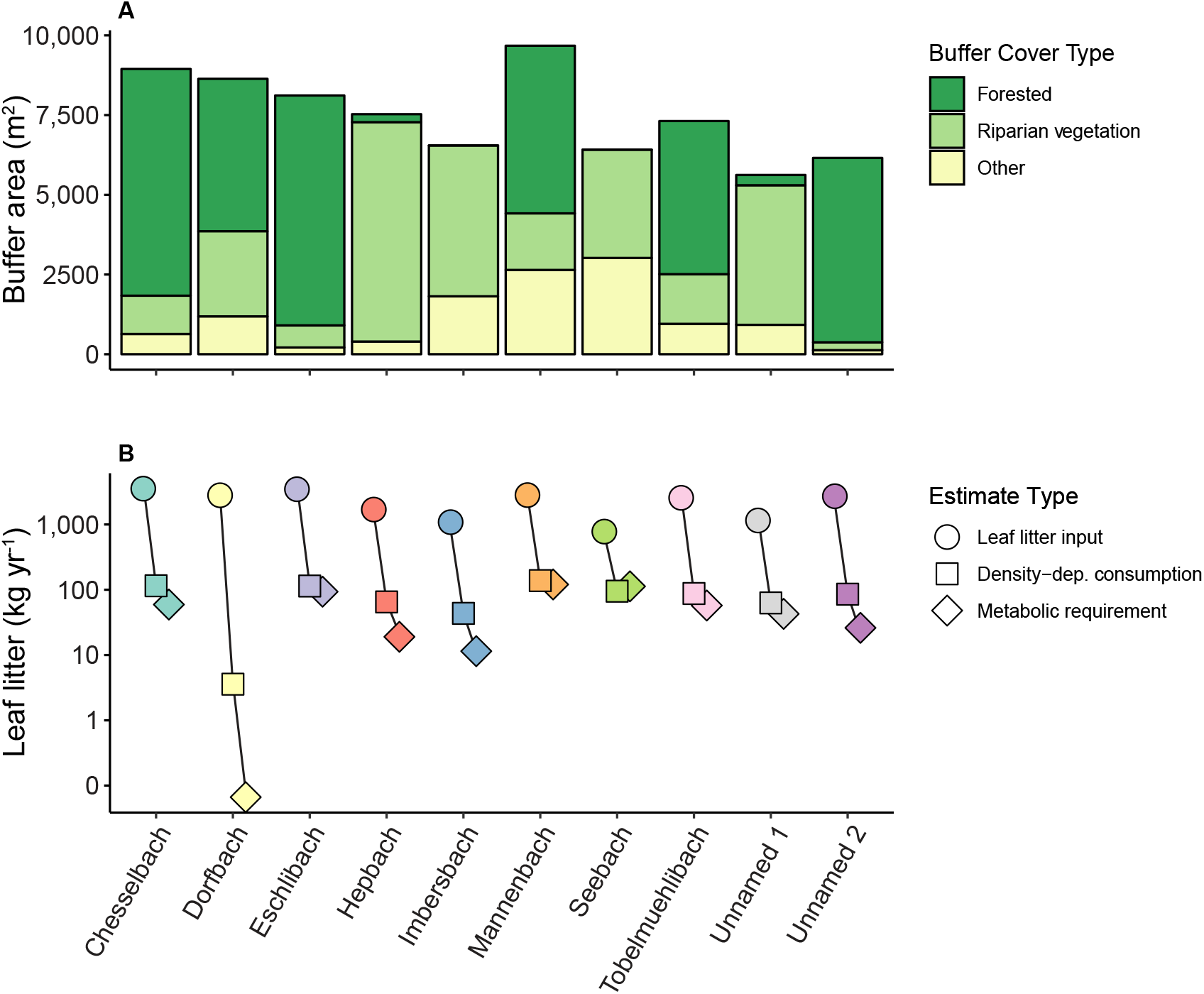
Leaf litter availability at the catchment scale. (A) Composition of the buffer area of each stream (one meter on either side), showing area of potential input from forest and other riparian vegetation. (B) Total potential leaf litter input in each catchment, based on composition of the buffer area in (A), compared to total leaf litter consumption from estimated amphipod abundance and density-dependent consumption rates (Table 1) and the estimated metabolic requirements of this same estimated population of amphipods. Note that y-axis is log-transformed in (B). The x-axis is the same for both panels.

## Discussion

As ecology moves towards a more predictive science, a central challenge is that the underlying mechanisms leading to an observed response – for example, ecosystem function – are occurring at a different scale (Levin 1992). In this context, the nonlinear relationships abundant in nature present challenges for upscaling. Often, it may be necessary to incorporate variance in the explanatory variable, not simply mean values, for predictions to be accurate: as Jensen’s inequality states, variance and skewness influence the integrals of nonlinear functions (Ruel and Ayres 1999; Martin and Huey 2008; Denny and Benedetti-Cecchi 2012). Using experimental manipulations at the level of individual small organisms, we found that local population density had a strong effect on leaf litter processing rates of two dominant freshwater detritivores, and thus, their per-capita (or per-biomass) contribution to ecosystem function. At the reach scale, the shape of this density-ecosystem function (DEF) relationship meant that estimated ecosystem function was similar across stream reaches, even when there was substantial spatial heterogeneity in organismal abundances. At the landscape scale, that is, the scale of riverine networks, the shape of this nonlinear relationship had strong implications for upscaled predictions of ecosystem function, because population density increases much faster than its corresponding ecosystem function. As a result, ecosystem function predictions based on our experimentally-derived DEF relationship were more than an order of magnitude lower than predictions made using more simplistic, mean-based estimates. Predictions based on DEF relationships also better aligned with the estimated consumption rate needed to meet the metabolic requirements of a catchment’s population. Thus, neglecting the role of density may systematically bias estimates of ecosystem function and lead to unrealistic predictions.

Intraspecific competition for resources is an essential regulator of population dynamics. Here, we demonstrate that intraspecific density could also regulate leaf litter processing, a key ecosystem function globally providing terrestrial resources to freshwater ecosystems (Tank et al. 2010; Gounand et al. 2018). Previously, leaf litter processing was shown to vary nonlinearly with abundance of macroinvertebrates, which was attributed to intraspecific competition for resources at high densities (Klemmer et al. 2012). However, in our experiments, resources were not limiting, and per-capita leaf processing decreased even at relatively low densities. Thus, we suggest two reasons for higher per-capita consumption at lower densities. First, we find it likely that interference competition (Schoener 1983) – competition for space rather than food resources (Moksnes 2004; Ward et al. 2007) – generated some of these nonlinearities. Secondly, our estimates of the metabolic requirements of different catchments’ populations suggest that individuals consume more than needed at low densities, while this interference competition reduces consumption rates to the metabolic minimum at high densities. In the broader context of upscaling, all mechanisms of intraspecific interaction, including exploitation and interference competition, are important as they could shape DEF relationships.

One main consequence of a nonlinear DEF relationship is that predictions at the landscape scale become challenging. This is especially relevant for organisms that are known to vary in their abundance locally over several orders of magnitude, such as the dominant shredders studied here. In our case, neglecting the role of density would lead to vast overestimates of ecosystem function; in other contexts (species, relationships, and functions) the reverse may be true. Connecting nonlinear population dynamics and spatial heterogeneity led to the development of scale transition theory (Melbourne and Chesson 2005; Chesson 2012), which has been applied to populations and communities and should be expanded to ecosystem-level processes. This nonlinear spatial averaging effect is likely to be of interest for optimizing regional ecosystem function. For example, extremely high densities of organisms may not provide the most “bang for their buck” in contributing to ecosystem function, since each additional individual contributes less and less; as such, some intermediate abundance may be optimal. In another perspective, this could be an argument for maintaining environmental heterogeneity and patches with high and low organismal abundance throughout a landscape. Even patches with relatively low density may have good ecosystem function, but source patches with high density are needed in order to supply colonists for sink patches which are more subject to demographic stochasticity, even if those high density patches do not necessarily have substantially higher ecosystem function themselves.

Our upscaling exercise relies on several simplifying assumptions. They include:

1. that the taxa in question are providing the bulk of the ecosystem function studied;
2. that the taxa have the same qualitative response to density in laboratory mesocosms as in natural habitats;
3. interspecific and trophic interactions do not additionally influence their per capita contribution to ecosystem function; and
4. that contributions to ecosystem function do not vary in space or time.

We justify the first assumption because, as discussed, the study taxa are numerically dominant among all macroinvertebrates in many European forested headwater streams, and their shredding function cannot easily be replaced by other taxa (Piscart et al. 2009; Nery and Schmera 2015). Furthermore, in these forested headwaters, our two lines of evidence suggested that abundance of these consumers, rather than leaf litter supply rates, should limit leaf litter processing (Figures 3 and 4). First, higher shredder abundances are not associated with higher leaf litter standing stock. Locally, leaf litter availability remained high even in some reaches with high densities of shredders. This was matched by our catchment-scale examination of leaf litter supply. Annual leaf litter inputs – estimated using several intentionally conservative assumptions – are one to several orders of magnitude greater than estimated consumption by spatially-heterogeneous shredder populations feeding according to density-dependent behavior.

Our measurements of vertical input during fall leaf were consistent with other published estimates of annual leaf litter input rates in forested and riparian shrub habitats in the Northern Hemisphere (reviewed in Conners and Naiman 2008). While there is spatiotemporal variation in the degree to which this total annual input is available to local communities in different part of the stream network, a substantial amount is stored in streams by debris dams, snags, and other features (Tank et al. 2010) and remains available for many months (Little and Altermatt 2018*c*). At the same time, leaf litter is transported downstream; while many inputs are retained near their point of entry to the stream, benthic leaf litter standing stock is not strongly driven by local vegetation characteristics and is rather redistributed through a stream network (Little and Altermatt 2018*c*), and thus available to consumers distant from its point of origin. Leaf litter inputs are more likely to be limiting in larger rivers, where inputs are an order of magnitude less per unit area (Naiman and Decamps 1997).

The other assumptions limit the quantitative realism of our upscaling exercise, but should not alter our main conclusions. For example, it is unlikely that the DEF relationship has the same parameters in natural streams with complex habitats as it does in simplified laboratory mesocosms. However, a negative exponential relationship would lead to the same qualitative effect of incorporating density-dependence on estimates of catchment-level ecosystem function, even if the shape of this relationship was somewhat different. Future work should consider how other mechanisms might modify this relationship. For example, we focused on a single trophic level of consumers, but interspecific interactions including competition and predation also influence behavior, consumption rates, and ecosystem function (Werner and Peacor 2003; Hines and Gessner 2012); a multi-trophic examination of the effects of dominant species density on ecosystem function is needed, including for cases unlike ours where resources are limiting and there is a functional response by consumers. It would also be of interest to investigate the assumption that the DEF relationship was the same in all patches, perhaps using simulation studies; habitat heterogeneity may alter organisms’ perception of density, and thus their behavior. Likewise, we doubt that temporal variation in the parameters of the DEF relationship would alter qualitative results, however simulations could explore how seasonal changes in water temperature and population age/size structure (Pöckl et al. 2003) would affect quantitative predictions of catchment-scale ecosystem function.

Our results expand the current understanding of biodiversity effects on ecosystem function (BEF) to include density-dependent effects on ecosystem function (DEF), recognizing that non-linear dependencies are prevalent and important (Grace et al. 2007; O’Connor et al. 2017). In fact, the two frameworks are related: release from intraspecific competition has been discussed as a mechanism through which increasing species richness accelerates ecosystem function (Jonsson and Malmqvist 2003; Weis et al. 2007; Patrick 2013). Connecting plot- and patch-level results to real, complex ecosystems and larger scales is recognized as one of the biggest challenges in ecosystem function research, with some debate as to the success of efforts to date (Hewitt et al. 2007; Snelgrove et al. 2014; Eisenhauer et al. 2016; Wardle 2016). Our results suggest that in some contexts, DEF may be extremely important: even without considering interactions with other species, intraspecific dynamics can be a strong control on ecosystem function, and thus accounting for organisms’ interactions will be essential in determining the drivers of ecosystem function.

## Acknowledgements

The authors thank two Samuel Hürlemann and Remo Wüthrich for help in the lab and field. CJL would like to thank the Eawag Eco PhD student writing group, organized by Heidi Kaech, for support and accountability; Joey Bernhardt and Simon Hart for valuable discussion; and Roman Alther and Isabelle Gounand for helpful comments on earlier manuscript drafts; and Volker Rudolf and two anonymous reviewers for comments on a later version. This is publication ISEM-YYYY-XXX of the Institut des Sciences de l’Evolution - Montpellier. Funding is from the Swiss National Science Foundation Grants PP00P3_150698 and PP00P3_179089 and the University of Zurich Research Priority Programme *URPP Global Change and Biodiversity* (to F.A.).

## Data Availability

Data will be made available on Dryad upon the manuscript’s acceptance.

## Supplementary Material

### Methods

#### Experimental details

The experiments were first performed on *Gammarus fossarum* in November 2016. We collected *G. fossarum* by kicknet from Sagentobelbach in Dübendorf, Switzerland (47.39° N, 8.59° E). Only adults were brought to the laboratory, where they were placed in large holding containers of ∼500 individuals, gradually brought up to 18 °C, and acclimated for two and a half days with ad libitum alder (*Alnus glutinosa* (Gaertner)) leaves conditioned for six days in stream water to establish natural microbial and fungal communities. In January 2017, we repeated the experiment with *D. villosus* individuals collected by kicknet from Lake Constance at Kesswil, Switzerland (47.60° N, 9.32° E).

For both focal species, we distributed individuals into experimental units so that medium and large individuals were equally represented in all replicates. With respect to sex, unless one divides individuals into males and females by separating precopulatory pairs, a microscope is necessary to identify sex. This is impractical in the field, and handling can injure the individuals. We instead assumed that subsequent allocation of individuals to treatments was random across the experiment.

**Figure S1.**
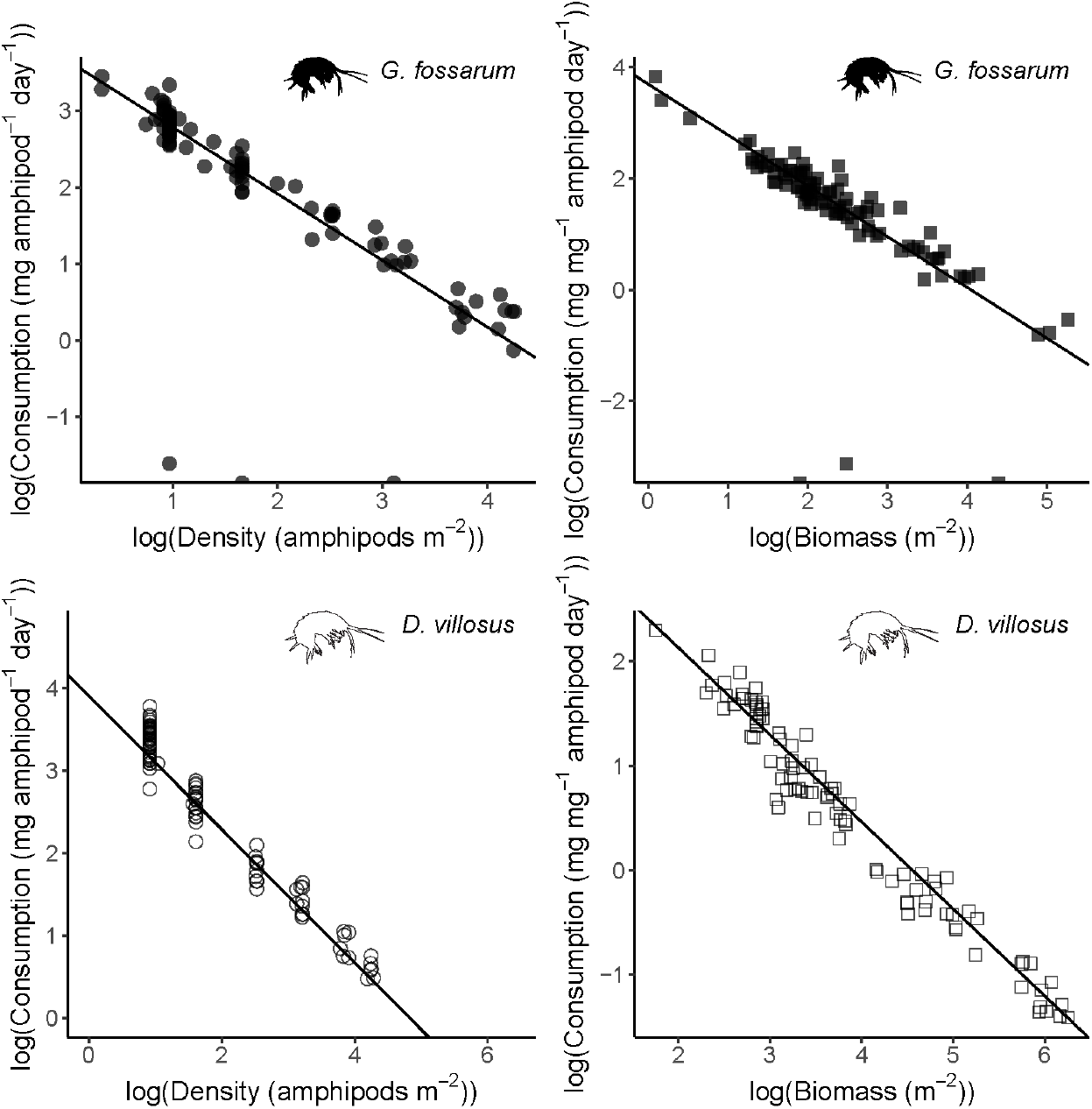
The association between density and per-capita consumption rate is linear in log-log space, for both G. fossarum and D. villosus, and both when measuring density as individual abundance or as biomass.

#### Data exploration and analysis for DEF relationships

Because the measures of density had long right tails, we explored density-consumption relationships which log-transformed this variable. We first used linear models to create (1) simple linear models, (2) linear models with log-transformed density data, and (3) linear models with log-transformed density and log-transformed consumption data. For this analysis, any zero values of consumption were replaced with 0.01 to enable log-transformation; fitting of nonlinear models (see below) used original data with zeros. To determine which type of model best fit the data, we examined diagnostic plots and tested each model’s residuals for normality using a Shapiro-Wilkes test. Non-normal residuals taken to indicate poor model fit, and we planned to use this initial comparison to determine which of three strategies to use for fitting the data: (1) if simple linear models had residuals that met assumptions of normality and homoscedasticity, we would use these models; (2) if the models with log-transformed density fit best we would re-fit the data using generalized linear models (GLM) with a log link function, and (3) if the third models fit best we would re-fit the data using non-linear least square models (NLS). However, we used normality of residuals in combination with other diagnostics to determine the best model for the data. Data were linear in log-log space for both species (Figure S1).

For *G. fossarum*, only the linear model (scenario 1 above) had normal residuals (Table S1). But diagnostics indicated that it did not fit the data well (Figure S2). While the residuals of the third model (on log-transformed density and consumption data) did not meet the assumption of normality according to the Shapiro-Wilkes test (Table S1), the other diagnostics indicate that the log-log model fit the data best other than three outliers where no leaf litter had been consumed (Figure S3). While deviating from normality, these observations did not have such high leverage (Cook’s distance, red lines in Figure S3) to exclude them, and would only make the nonlinear fit estimates more conservative by pulling down estimated consumption rates at low densities (Figure 1). Thus, we chose nonlinear model fits for both species.

**Table S1.** In order to determine the shape of the density-consumption relationship, residuals of different models (linear-linear, linear-log, and log-log) were tested for normality using a Shapiro-Wilkes test, where the null hypothesis is normality of the distribution. The G. fossarum dataset comprised 97 observations, and the D. villosus dataset 96 observations.

| **Model specification** | **Shapiro W statistic** | **Shapiro p-value** | **Model Adjusted R<sup>2</sup>** |
| --- | --- | --- | --- |
| *G. fossarum*, individual density |  |  |  |
| consumption ~ density | 0.98 | 0.09 | 0.45 |
| consumption ~ log(density) | 0.92 | < 0.001 | 0.73 |
| log(consumption) ~ log(density) | 0.31 | < 0.001 | 0.45 |
| *G. fossarum*, biomass density |  |  |  |
| consumption ~ density | 0.59 | < 0.001 | 0.13 |
| consumption ~ log(density) | 0.63 | < 0.001 | 0.52 |
| log(consumption) ~ log(density) | 0.37 | < 0.001 | 0.44 |
| *D. villosus*, individual density |  |  |  |
| consumption ~ density | 0.94 | < 0.001 | 0.46 |
| consumption ~ log(density) | 0.97 | 0.04 | 0.77 |
| log(consumption) ~ log(density) | 0.98 | 0.30 | 0.96 |
| *D. villosus*, biomass density |  |  |  |
| consumption ~ density | 0.90 | < 0.001 | 0.37 |
| consumption ~ log(density) | 0.94 | < 0.001 | 0.73 |
| log(consumption) ~ log(density) | 0.98 | 0.21 | 0.96 |

**Figure S2.**
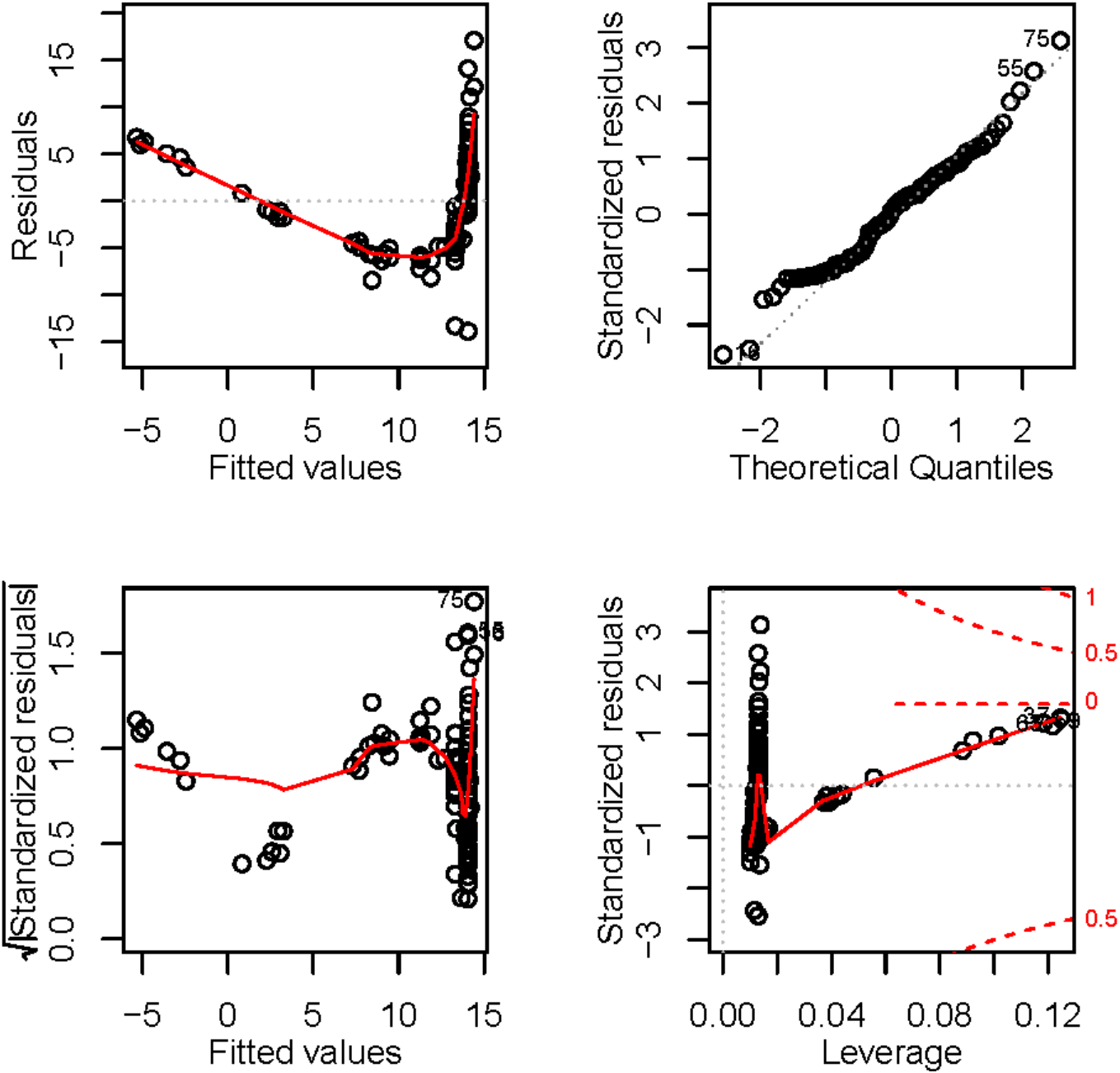
Diagnostic plots for the simple linear model of per-amphipod consumption rates as a function of density.

For the *D. villosus*, only the third linear model scenario (with log transformed density and consumption data) had normal residuals (see Table S1 above).

Therefore, we developed NLS models for both species. Parameter estimates from the linear models of transformed data were used as starting values for the NLS models. 95% confidence intervals for the model fits were generated by making 100,000 samples of coefficient estimates based on the approximate variance-covariance matrix extracted from the gNLS.

**Figure S3.**
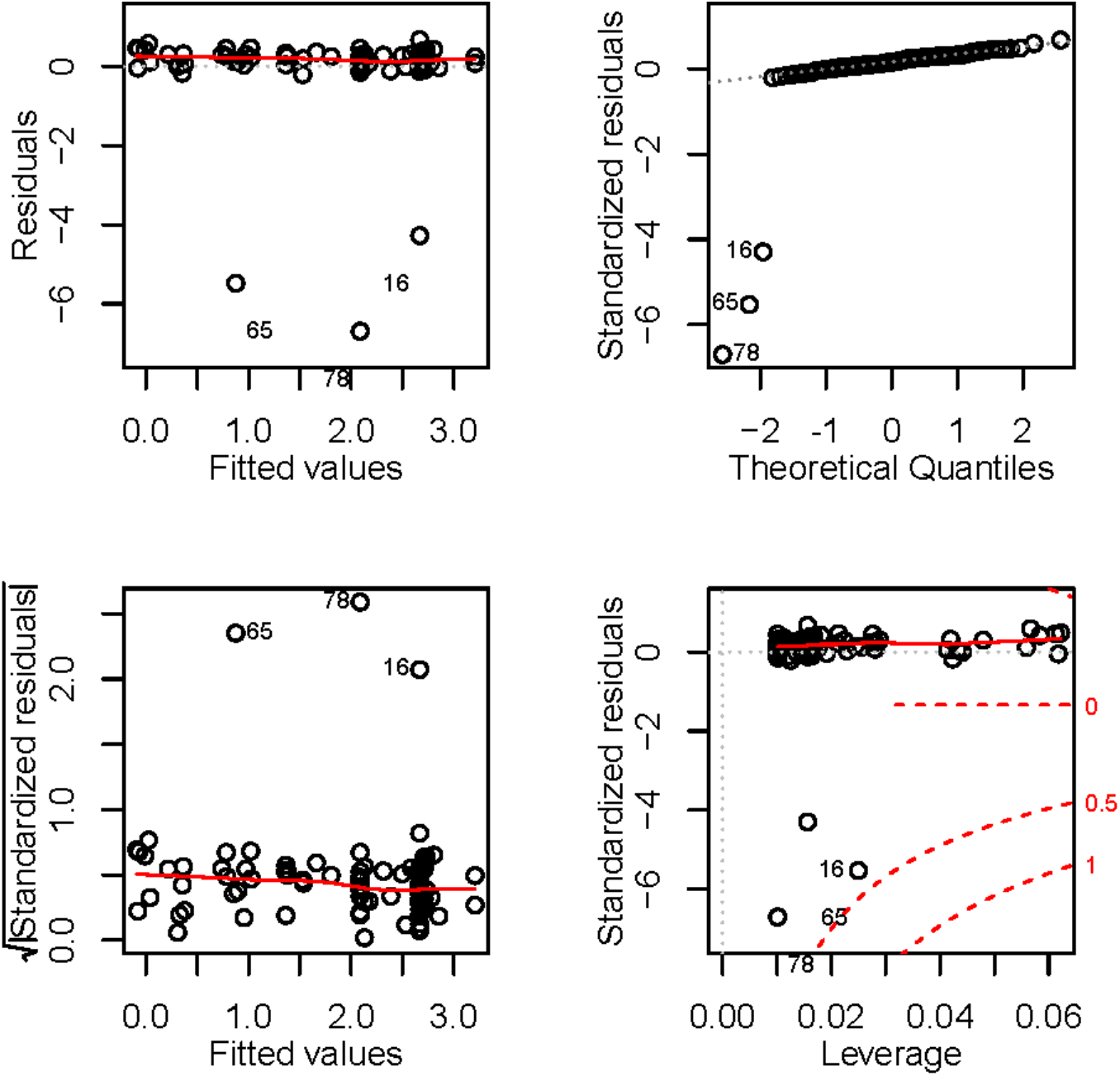
Diagnostic plots for the linear model of log-transformed per-amphipod consumption rates as a function of log-transformed density.

### Results

**Figure S4.**
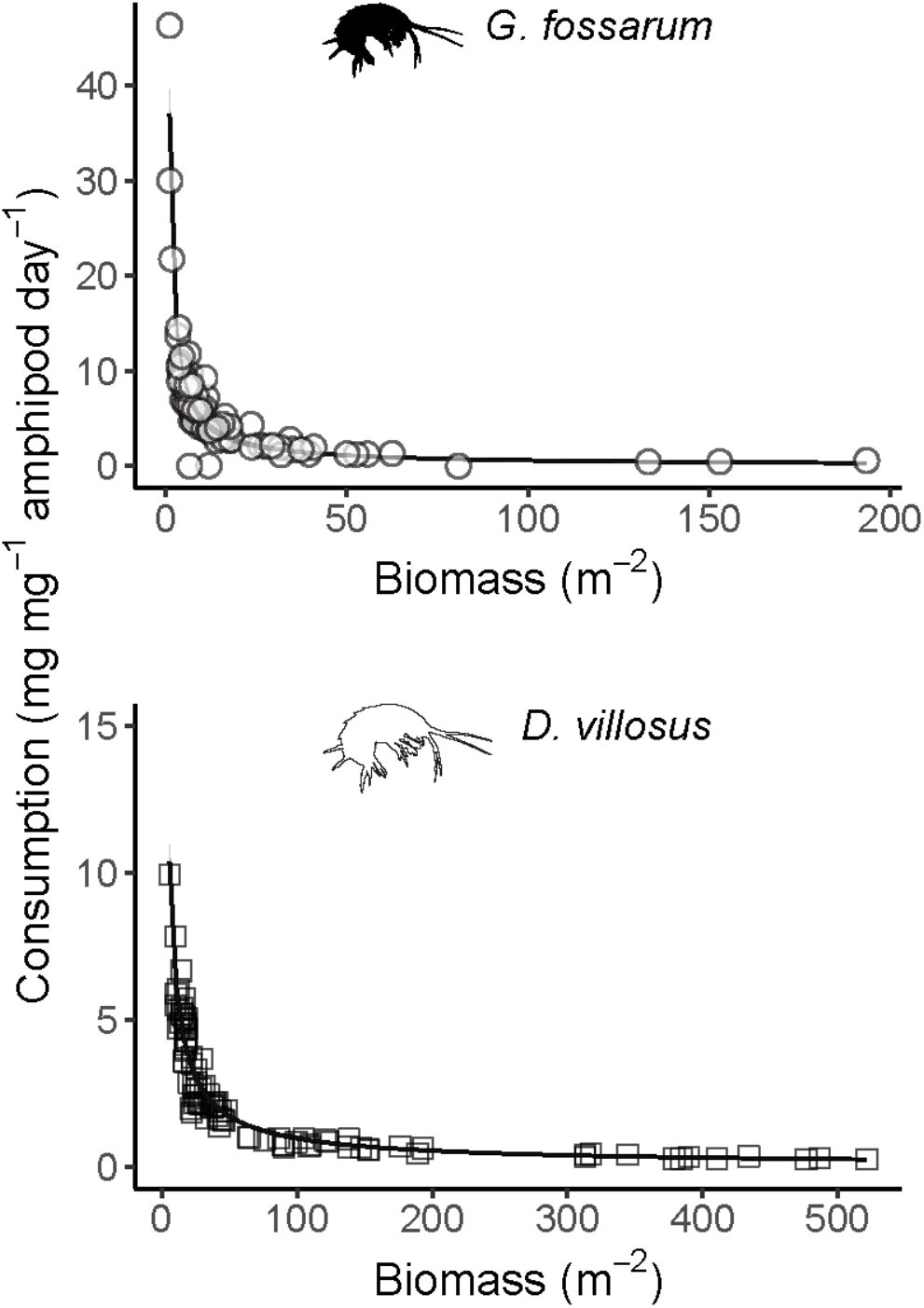
Negative exponential relationships between biomass and leaf litter consumption rates in experimental mesocosms for *G. fossarum* (consumption = 40.3*biomass^−0.92^) and *D. villosus* (consumption=44.7*biomass^−0.83^).

**Table S2.** Total amphipod abundance estimated using proportional estimation based on field sampling (*n* = number of sampling points in catchment, *l* = total stream length in the catchment), and three different upscaled estimates to whole-catchment leaf litter processing rates (grams/day). Briefly, Scenario 1 assumes density-independent per-capita processing; Scenario 2 assumes spatially-varying abundances in a catchment with leaf consumption calculated for each stream reach based on its particular abundance. Estimates are means of 1000 simulations, with 95% confidence intervals shown in parentheses.

| Stream name | Total Abundance | Scenario 1 Processing | Scenario 2 Processing |
| --- | --- | --- | --- |
| Chesselbach<br>$n = 13, l = 4.5 \text{ km}$ | 692,825<br>($\pm 1,035$) | 7,877.5<br>($\pm 11.8$) | 272.2<br>($\pm 0.1$) |
| Dorfbach<br>$n = 15, l = 4.3 \text{ km}$ | 1,587<br>($\pm 6$) | 18.0<br>($\pm 0.1$) | 13.6<br>($\pm 0.0$) |
| Eschlibach<br>$n = 12, l = 4.1 \text{ km}$ | 1,179,563<br>($\pm 1,249$) | 13,411.8<br>($\pm 14.2$) | 272.6<br>($\pm 0.1$) |
| Hepbach<br>$n = 13, l = 3.8 \text{ km}$ | 186,550<br>($\pm 622$) | 2,121.1<br>($\pm 7.1$) | 139.3<br>($\pm 0.1$) |
| Imbersbach<br>$n = 11, l = 3.3 \text{ km}$ | 166,930<br>($\pm 613$) | 1,898.0<br>($\pm 7.0$) | 53.7<br>($\pm 0.1$) |
| Mannenbach<br>$n = 15, l = 4.8 \text{ km}$ | 1,351,796<br>($\pm 1,408$) | 15,370.1<br>($\pm 16.0$) | 334.3<br>($\pm 0.1$) |
| Seebach<br>$n = 11, l = 3.2 \text{ km}$ | 1,434,449<br>($\pm 1,250$) | 16,309.9<br>($\pm 14.2$) | 250.1<br>($\pm 0.1$) |
| Tobelmühlibach<br>$n = 12, l = 3.7 \text{ km}$ | 738,789<br>($\pm 1,081$) | 8,400<br>($\pm 12.3$) | 183.4<br>($\pm 0.1$) |
| Unnamed Stream 1<br>$n = 9, l = 2.8 \text{ km}$ | 361,717<br>($\pm 849$) | 4,112.8<br>($\pm 9.6$) | 74.0<br>($\pm 0.1$) |
| Unnamed Stream 2<br>$n = 10, l = 3.6 \text{ km}$ | 322,401<br>($\pm 672$) | 3,665.7<br>($\pm 7.6$) | 220.1<br>($\pm 0.0$) |

**Figure S5.**
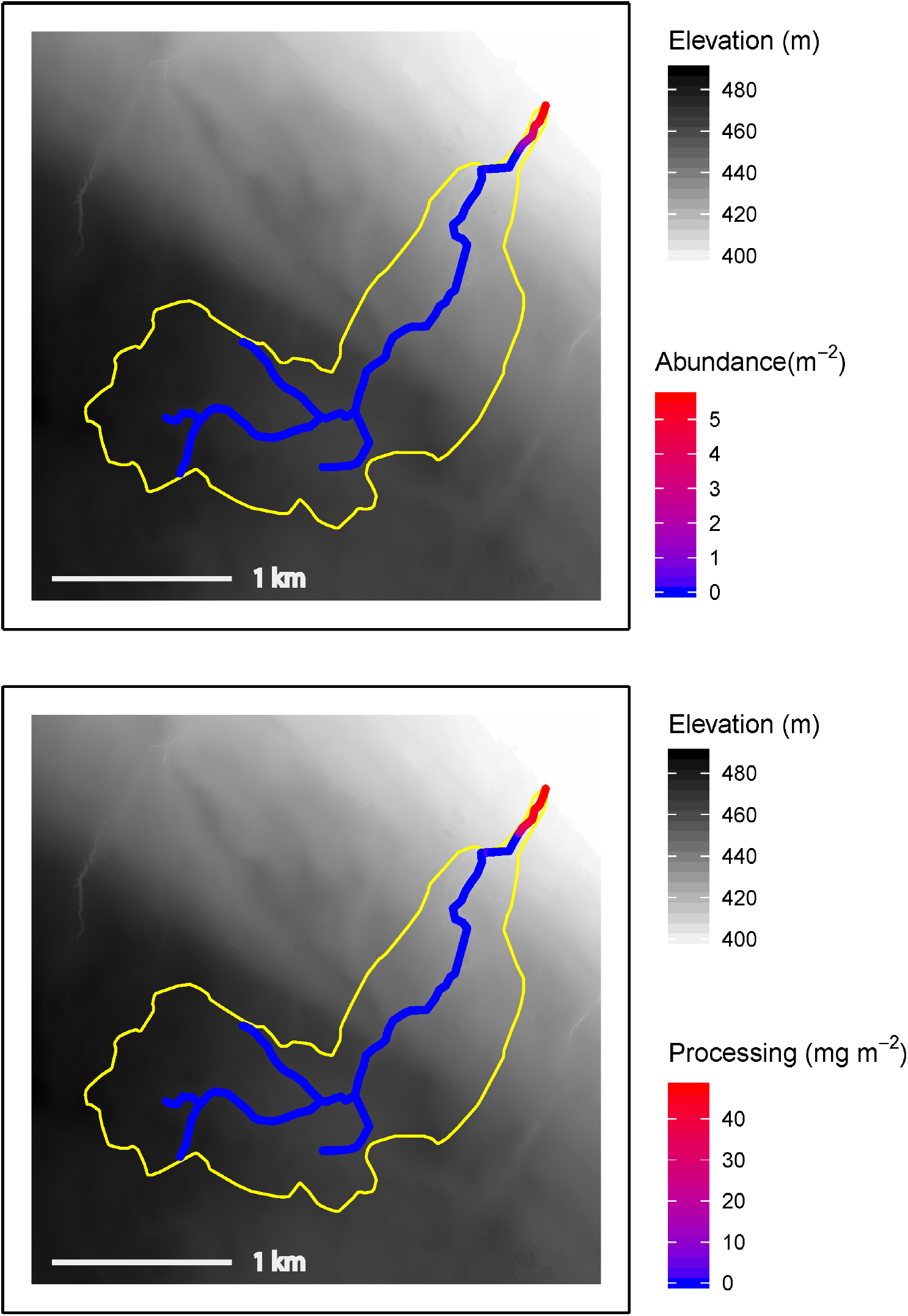
Hotspots of abundance (top panel) and leaf litter processing (bottom panel) in the Dorfbach stream catchment (outlines in yellow), based on upscaling abundance data from 15 sampling points up to longitudinal abundance distributions using inverse distance weighted interpolation. Processing rates (per day) were calculated by multiplying the interpolated abundance in a 1 m section of stream length by the experimentally-derived negative power relating *G. fossarum* density to per-capita leaf litter consumption. This figure shows the mean of 1000 simulations of the interpolation process. Data sources: swisstopo (2010, 2014), Vector25 and TLM3D, DV 5704 000 000, reproduced by permission of swisstopo/JA100119.

**Figure S6.**
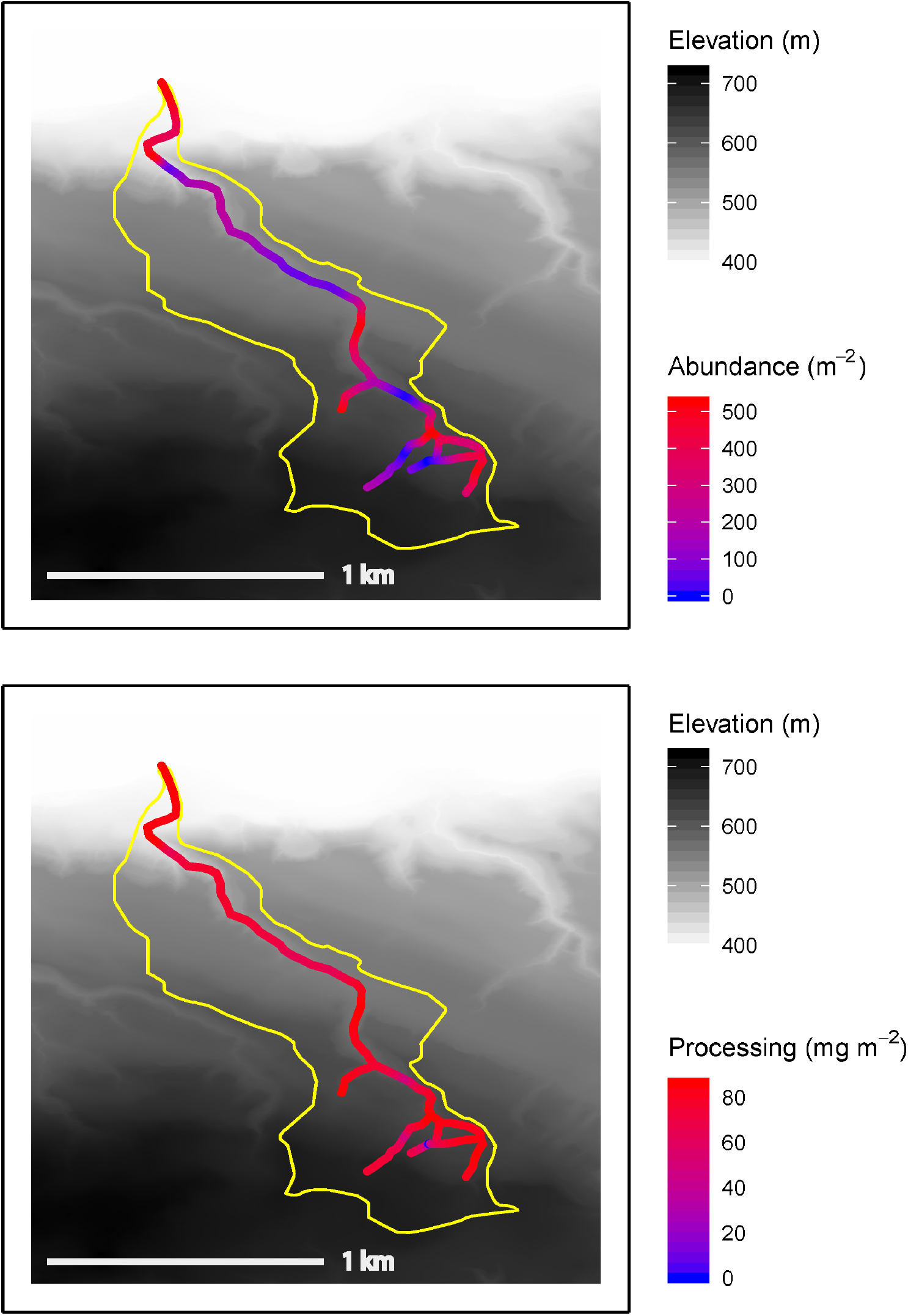
Hotspots of abundance (top panel) and leaf litter processing (bottom panel) in the Eschlibach stream catchment (outlines in yellow), based on upscaling abundance data from 12 sampling points up to longitudinal abundance distributions using inverse distance weighted interpolation. Processing rates (per day) were calculated by multiplying the interpolated abundance in a 1 m section of stream length by the experimentally-derived negative power relating *G. fossarum* density to per-capita leaf litter consumption. This figure shows the mean of 1000 simulations of the interpolation process. Data sources: swisstopo (2010, 2014), Vector25 and TLM3D, DV 5704 000 000, reproduced by permission of swisstopo/JA100119.

**Figure S7.**
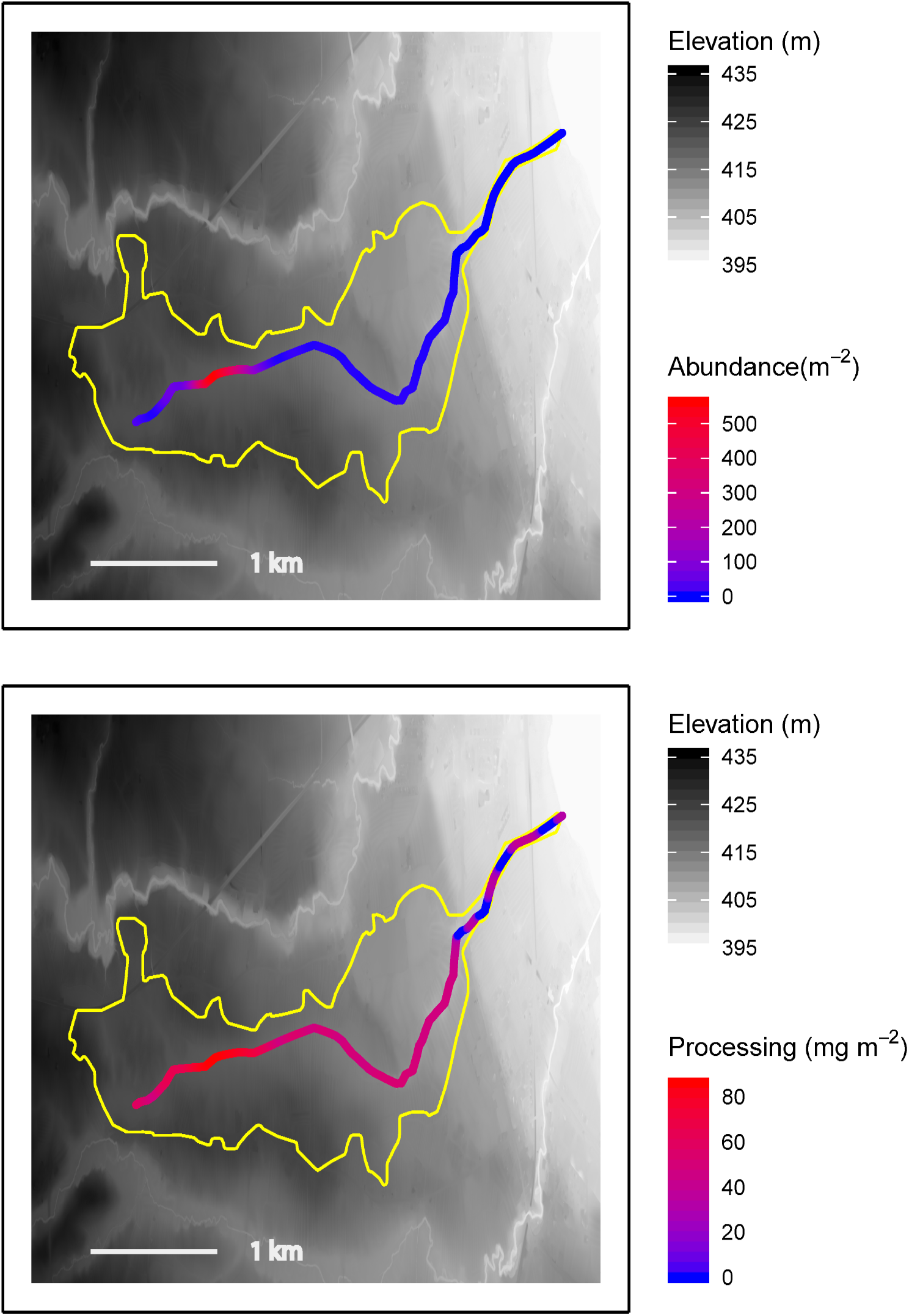
Hotspots of abundance (top panel) and leaf litter processing (bottom panel) in the Hepbach stream catchment (outlines in yellow), based on upscaling abundance data from 13 sampling points up to longitudinal abundance distributions using inverse distance weighted interpolation. Processing rates (per day) were calculated by multiplying the interpolated abundance in a 1 m section of stream length by the experimentally-derived negative power relating *G. fossarum* density to per-capita leaf litter consumption. This figure shows the mean of 1000 simulations of the interpolation process. Data sources: swisstopo (2010, 2014), Vector25 and TLM3D, DV 5704 000 000, reproduced by permission of swisstopo/JA100119.

**Figure S8.**
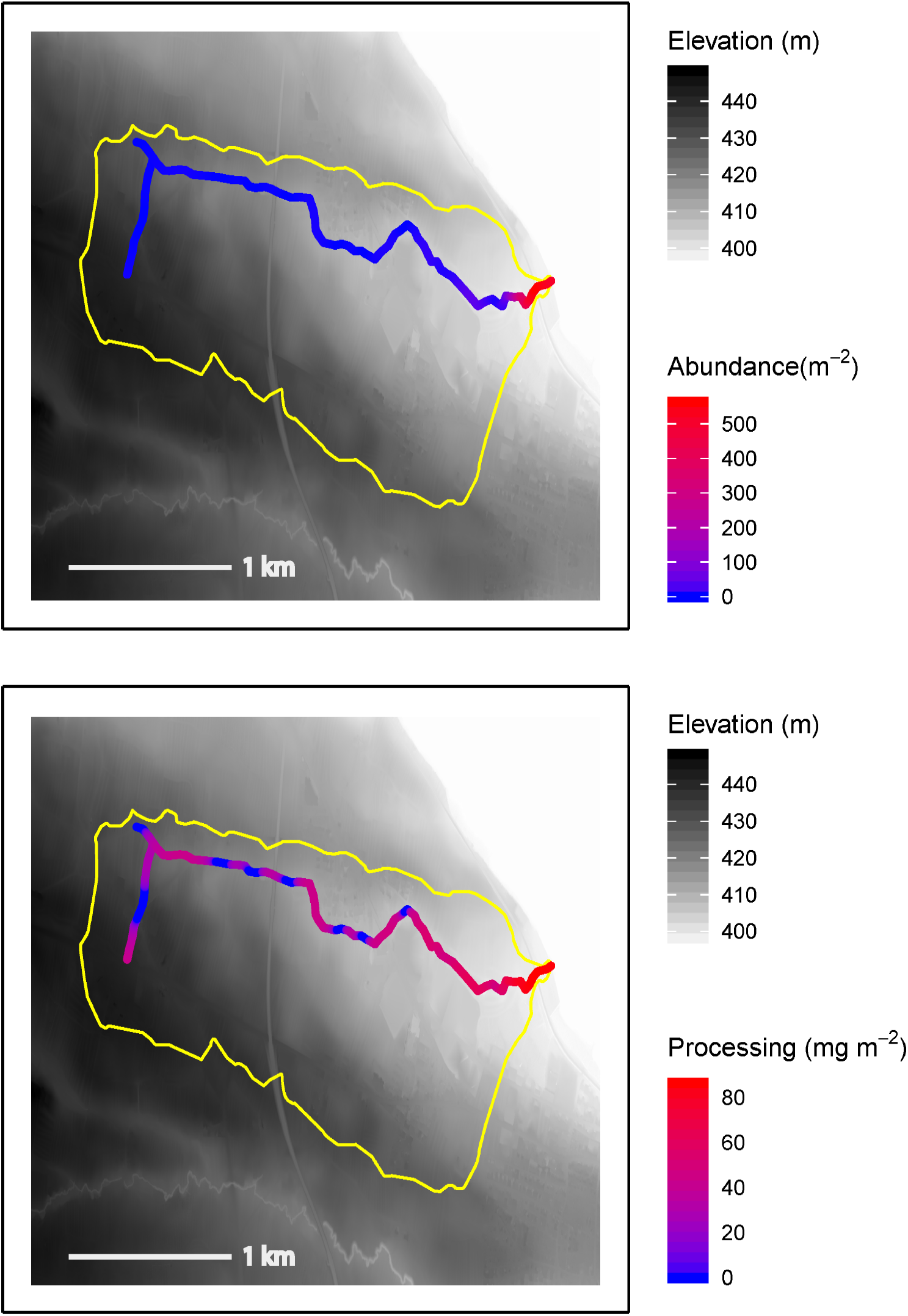
Hotspots of abundance (top panel) and leaf litter processing (bottom panel) in the Imbersbach stream catchment (outlines in yellow), based on upscaling abundance data from 11 sampling points up to longitudinal abundance distributions using inverse distance weighted interpolation. Processing rates (per day) were calculated by multiplying the interpolated abundance in a 1 m section of stream length by the experimentally-derived negative power relating *G. fossarum* density to per-capita leaf litter consumption. This figure shows the mean of 1000 simulations of the interpolation process. Data sources: swisstopo (2010, 2014), Vector25 and TLM3D, DV 5704 000 000, reproduced by permission of swisstopo/JA100119.

**Figure S9.**
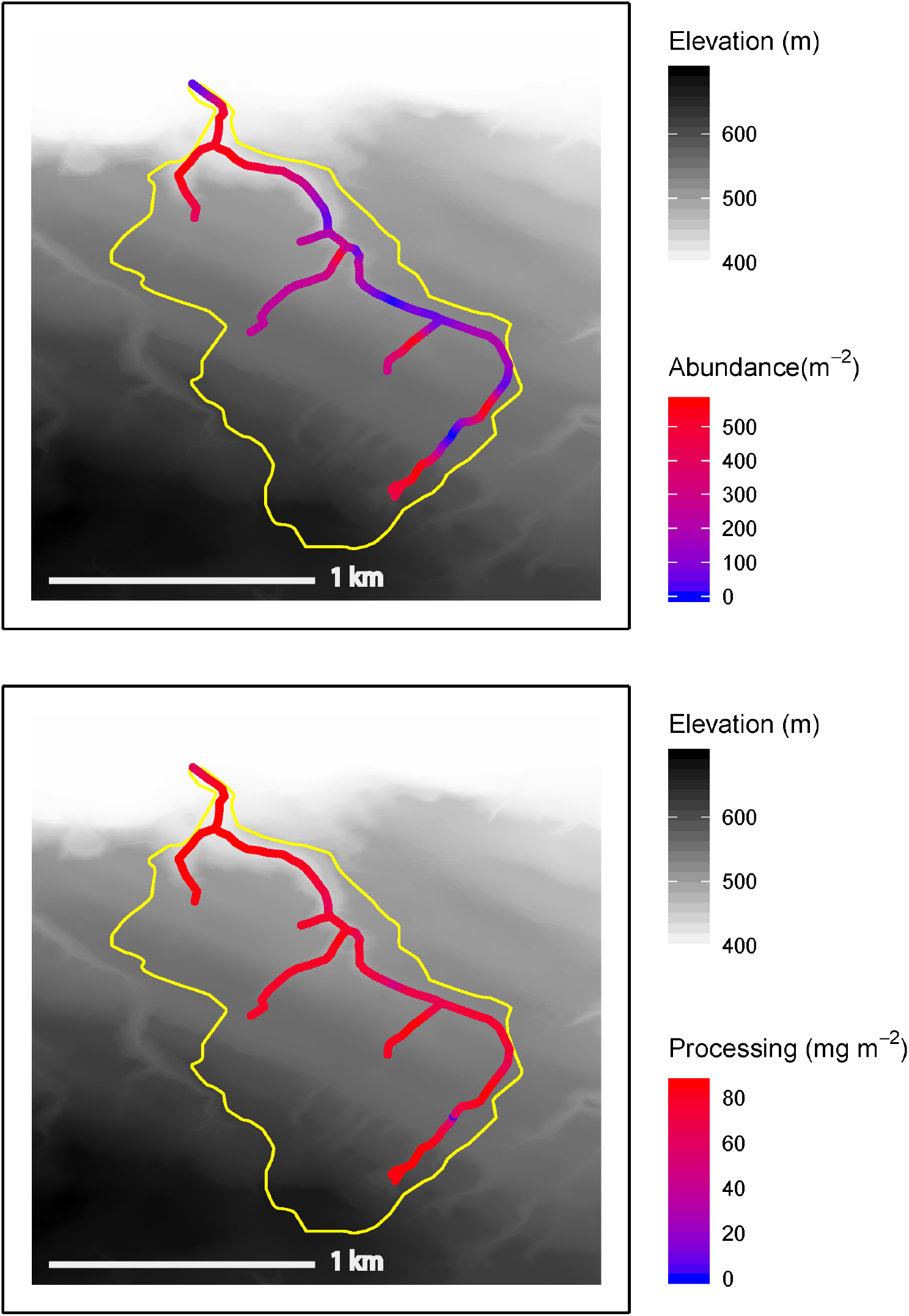
Hotspots of abundance (top panel) and leaf litter processing (bottom panel) in the Mannenbach stream catchment (outlines in yellow), based on upscaling abundance data from 15 sampling points up to longitudinal abundance distributions using inverse distance weighted interpolation. Processing rates (per day) were calculated by multiplying the interpolated abundance in a 1 m section of stream length by the experimentally-derived negative power relating *G. fossarum* density to per-capita leaf litter consumption. This figure shows the mean of 1000 simulations of the interpolation process. Data sources: swisstopo (2010, 2014), Vector25 and TLM3D, DV 5704 000 000, reproduced by permission of swisstopo/JA100119.

**Figure S10.**
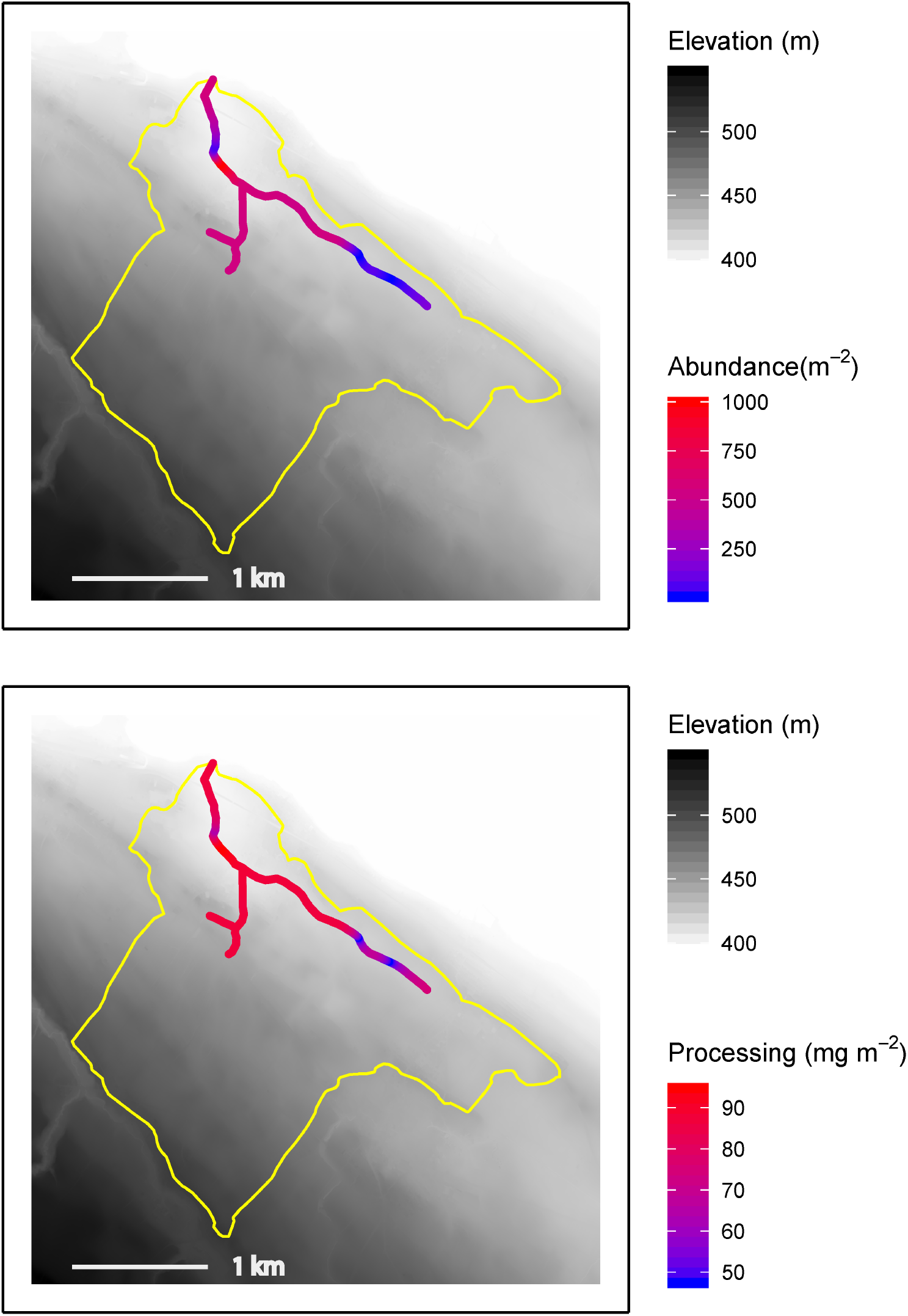
Hotspots of abundance (top panel) and leaf litter processing (bottom panel) in the Seebach stream catchment (outlines in yellow), based on upscaling abundance data from 11 sampling points up to longitudinal abundance distributions using inverse distance weighted interpolation. Processing rates (per day) were calculated by multiplying the interpolated abundance in a 1 m section of stream length by the experimentally-derived negative power relating *G. fossarum* density to per-capita leaf litter consumption. This figure shows the mean of 1000 simulations of the interpolation process. Data sources: swisstopo (2010, 2014), Vector25 and TLM3D, DV 5704 000 000, reproduced by permission of swisstopo/JA100119.

**Figure S11.**
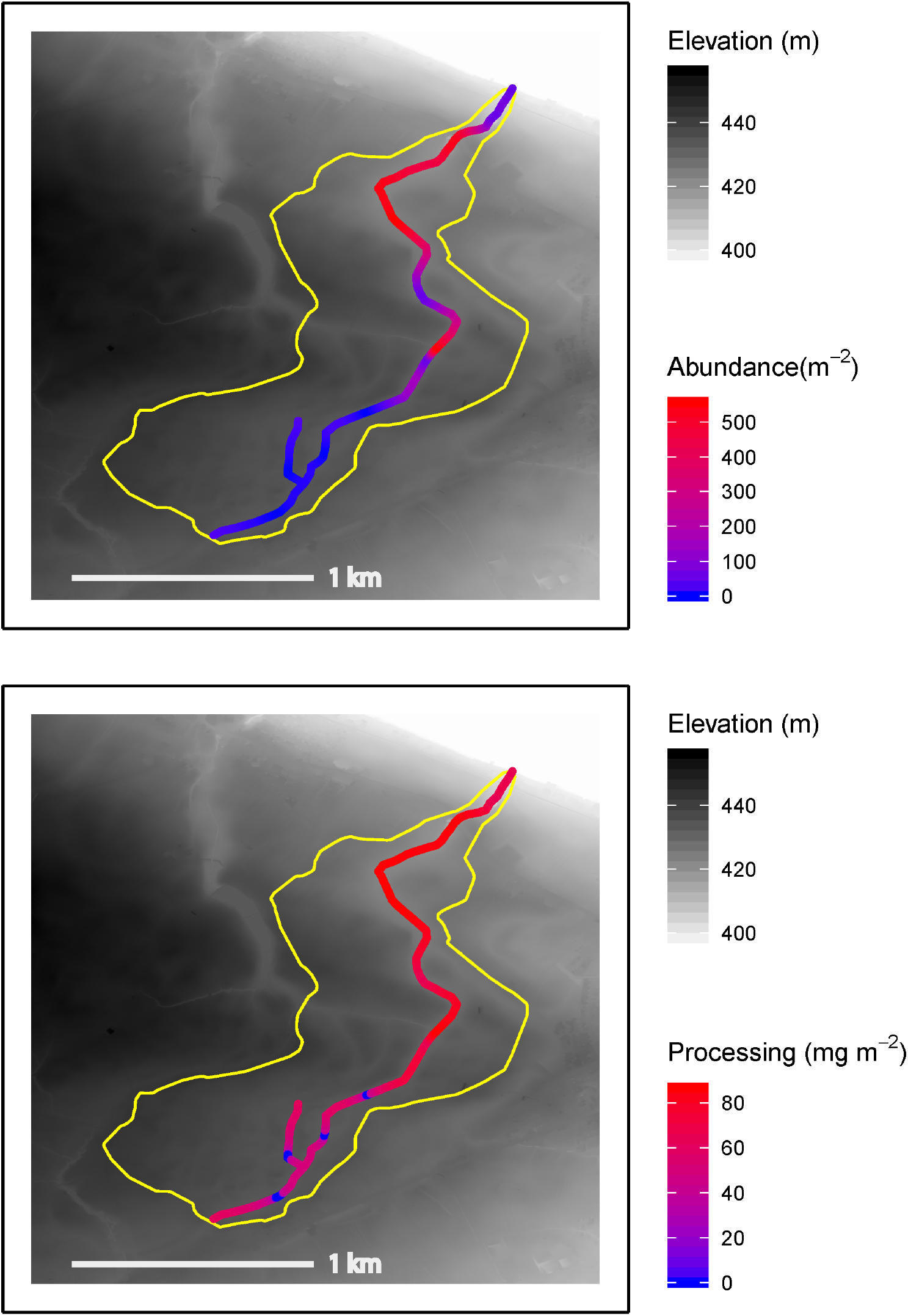
Hotspots of abundance (top panel) and leaf litter processing (bottom panel) in the Tobelmühlibach stream catchment (outlines in yellow), based on upscaling abundance data from 12 sampling points up to longitudinal abundance distributions using inverse distance weighted interpolation. Processing rates (per day) were calculated by multiplying the interpolated abundance in a 1 m section of stream length by the experimentally-derived negative power relating *G. fossarum* density to per-capita leaf litter consumption. This figure shows the mean of 1000 simulations of the interpolation process. Data sources: swisstopo (2010, 2014), Vector25 and TLM3D, DV 5704 000 000, reproduced by permission of swisstopo/JA100119.

**Figure S12.**
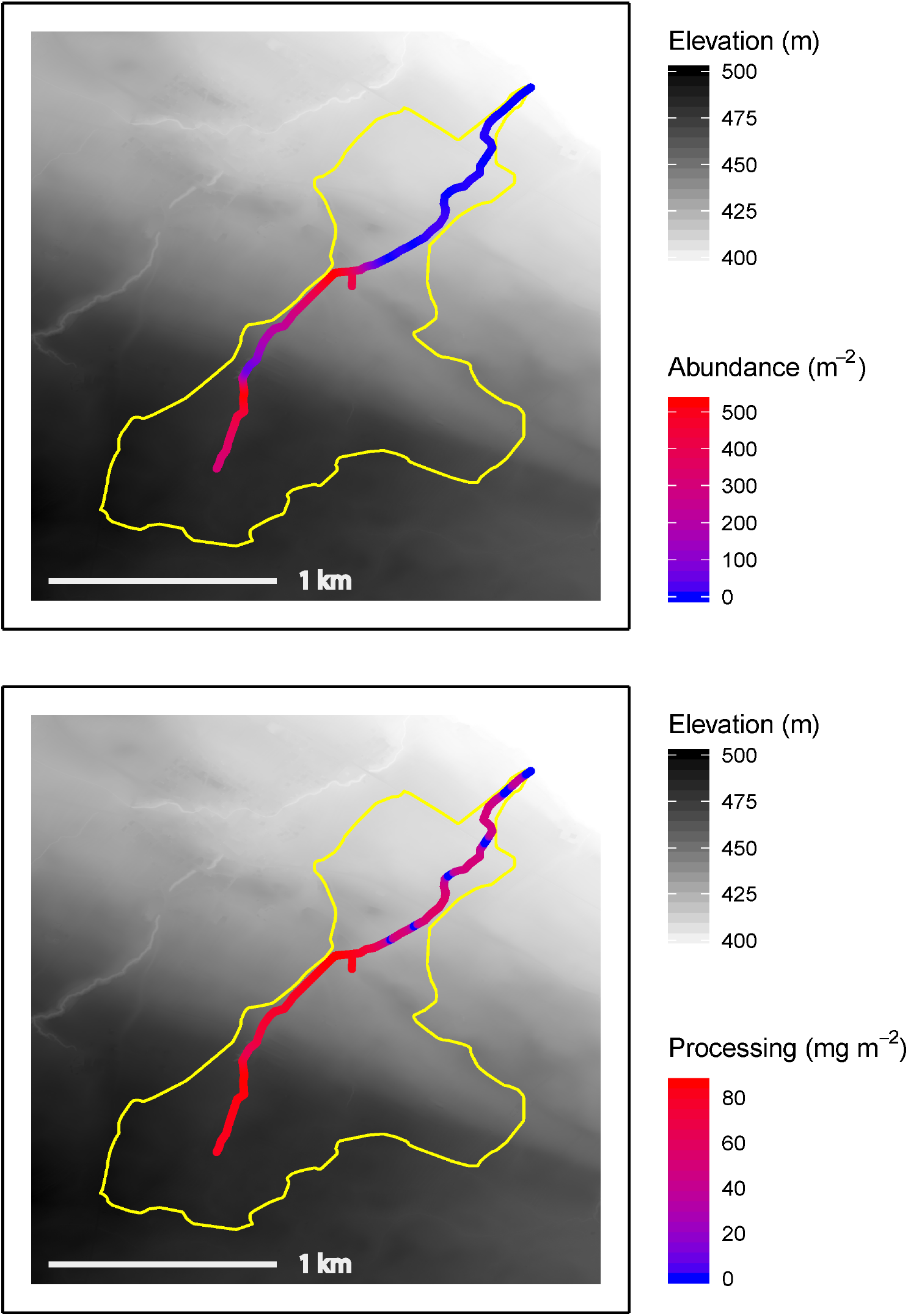
Hotspots of abundance (top panel) and leaf litter processing (bottom panel) in the Unnamed Stream #1 catchment (outlines in yellow), based on upscaling abundance data from 9 sampling points up to longitudinal abundance distributions using inverse distance weighted interpolation. Processing rates (per day) were calculated by multiplying the interpolated abundance in a 1 m section of stream length by the experimentally-derived negative power relating *G. fossarum* density to per-capita leaf litter consumption. This figure shows the mean of 1000 simulations of the interpolation process. Data sources: swisstopo (2010, 2014), Vector25 and TLM3D, DV 5704 000 000, reproduced by permission of swisstopo/JA100119.

**Figure S13.**
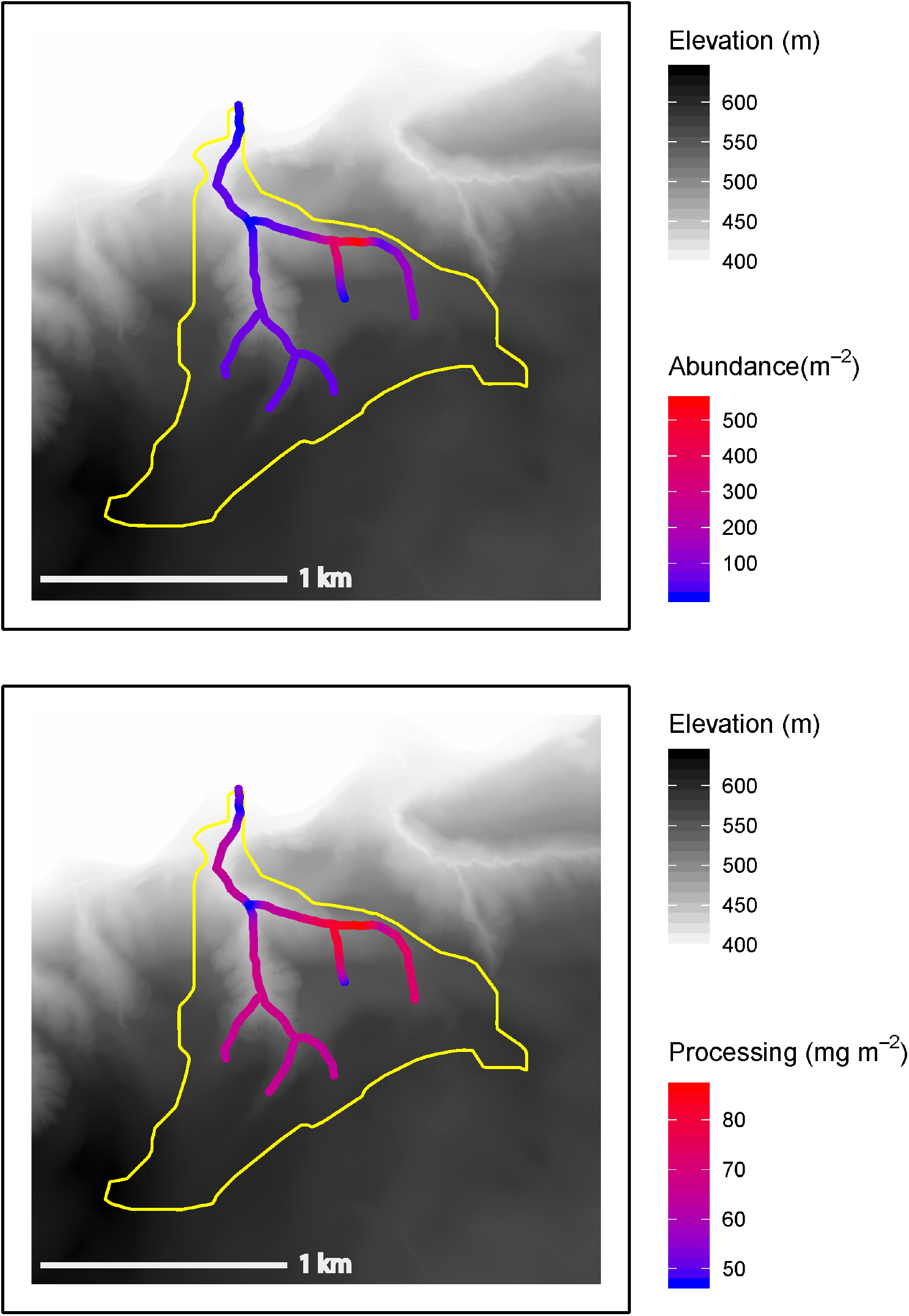
Hotspots of abundance (top panel) and leaf litter processing (bottom panel) in the Unnamed Stream #2 catchment (outlines in yellow), based on upscaling abundance data from 10 sampling points up to longitudinal abundance distributions using inverse distance weighted interpolation. Processing rates (per day) were calculated by multiplying the interpolated abundance in a 1 m section of stream length by the experimentally-derived negative power relating *G. fossarum* density to per-capita leaf litter consumption. This figure shows the mean of 1000 simulations of the interpolation process. Data sources: swisstopo (2010, 2014), Vector25 and TLM3D, DV 5704 000 000, reproduced by permission of swisstopo/JA100119.

**Figure S14.**
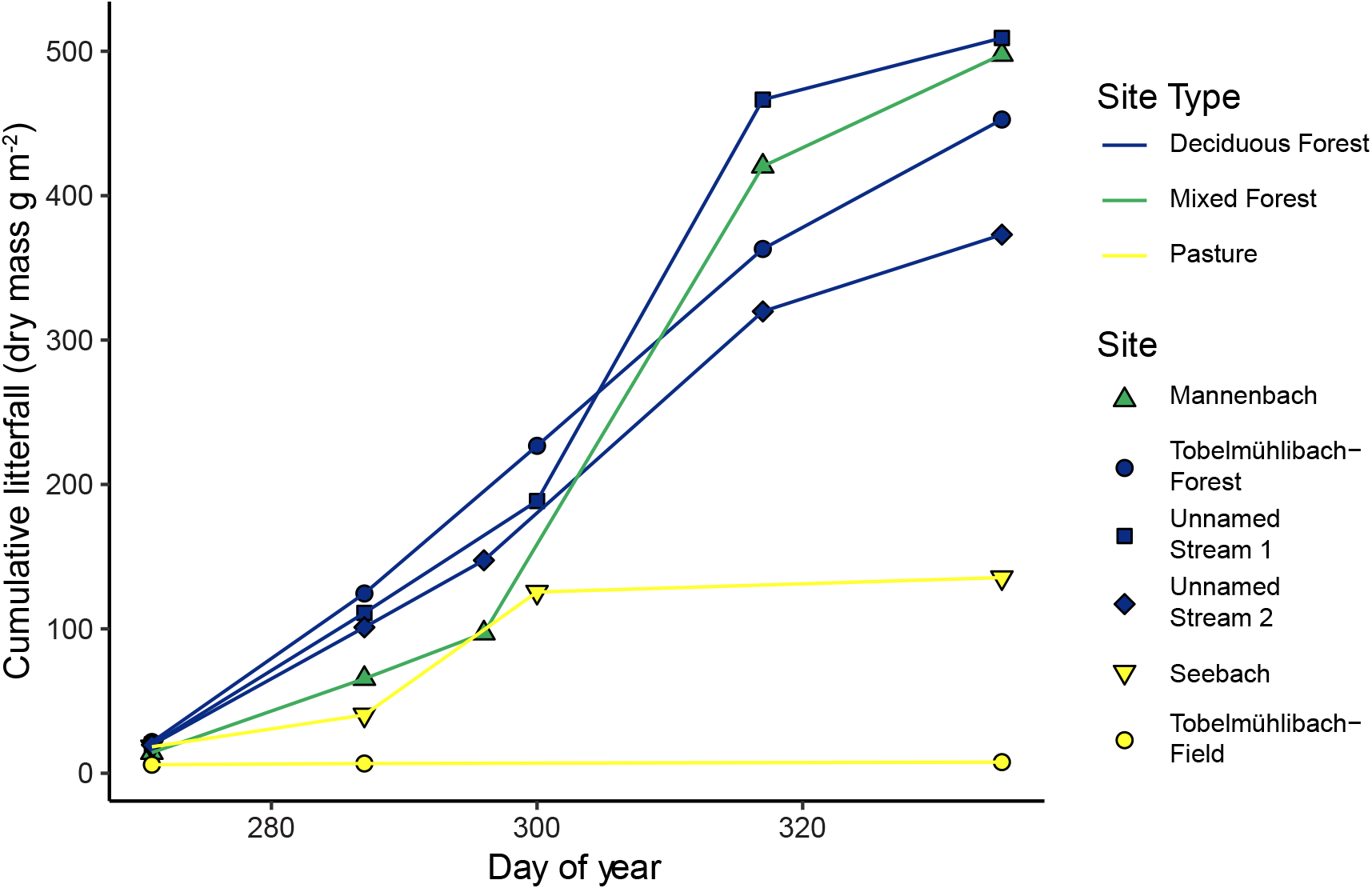
Accumulation of leaf litter through the period of autumn leaf drop at four different locations in the study area. At each location, eight litter traps of 800 square centimeters each were deployed. Traps in forested areas were emptied every nine to 18 days and traps in pasture areas were emptied less frequently because less litter accrued in them. Litterfall in a mixed pine/deciduous forest site was similar (498 g m-2 dry weight) to that of the three deciduous forest sites (453, 509 and 373 g m-2 dry weight), so all four sites were averaged to produce our estimate of annual litter input in forested areas.

